# Novel integrated risk index reveals uneven impact of climate change on fascioliasis risk across Africa and Europe

**DOI:** 10.64898/2025.12.04.692276

**Authors:** Tiem van der Deure, Samson Mukaratirwa, Moses Chimbari, Agrippa Dube, Tawanda Manyangadze, Mita Eva Sengupta, Safari Kinunghi, Rubens Mita, Julia Walker, Bryan O. Nyawanda, David Nogués-Bravo, Anna-Sofie Stensgaard

**Affiliations:** Department of Veterinary and Animal Sciences, University of Copenhagen, Copenhagen, Denmark; Center for Macroecology, Evolution, and Climate, Globe Institute, University of Copenhagen, Copenhagen, Denmark; School of Agriculture and Science, College of Agriculture, Engineering and Science, University of KwaZulu-Natal, Durban, South-Africa; One Health Center for Zoonoses and Tropical Veterinary Medicine, Ross University School of Veterinary Medicine, Basseterre, Saint Kitts and Nevis; School of Medicine, College of Health Sciences, University of KwaZulu-Natal, Durban, South Africa; Department of Physics, Geography and Environmental Sciences, School of Natural Sciences, Great Zimbabwe University, Masvingo, Zimbabwe; Geosciences Department, School Geosciences, Disaster and Sustainable Development, Faculty of Science, Bindura University of Science and Education, Bindura, Zimbabwe; National Institute for Medical Research (NIMR), Mwanza Centre, Mwanza, Tanzania; Swiss Tropical and Public Health Institute, Allschwil, Switzerland

**Author notes:** Corresponding authors: Tiem van der Deure, and Anna-Sofie Stensgaard.

## Abstract

Fascioliasis, a zoonotic parasitic disease caused by the liver flukes *Fasciola hepatica* and *Fasciola gigantica*, poses a significant threat to livestock and human health globally. Climate change is altering the transmission dynamics of this disease; however, the direction and magnitude of this change remain uncertain due to a lack of mechanistically grounded predictions at continental scale. In this study, we developed a novel risk index to assess how climate change may affect the potential for fascioliasis transmission across Africa and Europe. The index integrates the following three components that together capture the environmental requirements for *Fasciola* transmission: the temperature dependence of the parasite transmission; climatic suitability for the snail intermediate host; and availability of freshwater habitats. To estimate these components, we used data from published experimental studies, thousands of snail occurrence records, and both mechanistic and correlative modelling approaches.

We predict that climate change will generally reduce fascioliasis risk across most of the ranges of the two investigated liver fluke species, albeit with marked geographical differences. The transmission risk of *F. gigantica* is predicted to decrease in the Sahel and West Africa, due to above-optimal temperatures for parasite transmission and climatic suitability for the intermediate hosts. However, risk for *F. hepatica* transmission is projected to increase in parts of Northern Europe, Scandinavia, and Iceland, where livestock densities are high. We also find that the potential for hybridization between *F. hepatica* and *F. gigantica* may decline due to reduced geographic overlap in areas that are highly suitable for transmission.

By providing a scalable, ecologically informed framework for predicting fascioliasis risk under climate change, this study lays the groundwork for improved local risk assessment. Our results provide a more nuanced perspective on the potential impact of climate change on fascioliasis transmission, showing that this will be highly uneven across Africa and Europe.

**Author summary:** Fascioliasis is a snail-borne, zoonotic disease that affects both humans and livestock, causing major health burden and economic losses worldwide. The liver flukes that cause the disease use freshwater snails as their intermediate hosts. They therefore require aquatic habitats and suitable temperatures to complete their life cycle. Climate change altering these conditions; however, it is unclear how this will affect transmission of liver flukes in the future. Here we develop a new risk index that combines three key environmental factors of importance for liver fluke transmission: how suitable temperatures are for parasite transmission, the climatic suitability for snail hosts, and the availability of water. We mapped current and projected future transmission risk for two main species, *Fasciola hepatica* and *Fasciola gigantica*, across Africa and Europe. Different from what has been predicted by earlier studies, our results suggest that transmission risk will decline in across most of the range of both species. This will have particularly big impacts in areas where livestock densities are high, including the Sahel region. In northern Europe, transmission risk increases slightly. This study also provides a modelling framework and novel ecological knowledge that can be used to assess risk of transmission at local and continental scale.

## 1 Introduction

Fascioliasis is a re-emerging, zoonotic, parasitic disease with a global distribution that causes significant burden on animal and human health [1]. The disease is caused by infection with liver flukes of the genus *Fasciola*. It is one of the main causes of food-borne trematodiasis, with an estimated 2.6 million people infected worldwide [2–5]. In livestock, prevalences of 50% or higher have been reported [6], severely impacting animal welfare and agricultural productivity [7]. The two most important *Fasciola* species are *Fasciola hepatica*, which is found on all continents but most common in temperate areas, and *Fasciola gigantica*, which is restricted to tropical areas [8]. These species are able to hybridize, and hybrid forms have been linked to the emergence of fascioliasis in some areas [9,10].

*Fasciola* parasites have a complex life cycle which includes alternating between free-living stages, stages in a mammalian definitive host, and stages in a freshwater snail intermediate host. The rates of development of liver fluke eggs and the intramolluscan stages of the parasite increase with higher temperatures and cease below a certain temperature threshold [11], making liver fluke transmission highly sensitive to global warming. Likewise, liver fluke development is dependent on the presence of competent intermediate snail hosts, which are also highly sensitive to changes in environmental conditions, including the availability of aquatic habitats [12,13]. These freshwater habitats are also important for the parasite directly, as liver fluke eggs hatch in water and cercariae encyst and attach to aquatic vegetation. Consistently, a major study found that temperature and rainfall are key predictors for the prevalence of *F. hepatica* in European livestock [14].

Elucidating how climate change could affect the dynamics of *Fasciola* transmission globally is key to preparedness in the face of the re-emergence of fascioliasis [15,16], and several approaches have been used to address this challenging question. Firstly, multiple studies have used climate model predictions of temperature and precipitation to create future projections of the Ollerenshaw index [17], a monthly risk index developed to advise farmers in Wales, or adaptations to it [18–22]. For instance, Fox et al. [23] and Caminade et al. [24] applied this index to the United Kingdom and Europe, respectively, and predicted that fascioliasis risk has already increased since the 1970s and will continue to increase. Another approach is to use statistical models to find correlations between the observed prevalence of *Fasciola* infection and environmental variables [25,26]. Using such an approach, Cuervo et al. connected increasing temperatures to occurrence of *Fasciola* and its host snails at increasingly high altitudes in northern Bolivia, where human fascioliasis is hyperendemic [27]. Finally, a third approach is to use a mechanistic model of the entire *Fasciola* life cycle [28,29]. Using such a model, Smith [28] found that the number of metacercariae in Great Britain would increase due to climate change.

While these studies generated important insights on the impact of climate change on fascioliasis, they have limitations. The Ollerenshaw index has limited predictive power [30] and was not designed for continental-scale modelling or predicting climate change impacts. Correlative models can yield great insight into drivers of transmission where occurrence data availability is sufficient; however, fascioliasis has not been comprehensively mapped, with data lacking particularly for animal infections and especially in Africa. Similarly, mechanistic models have high data input requirements, including data on snail traits and local hydrology, and have thus far only been applied on local scales [29]. It is therefore highly uncertain how climate change will change fascioliasis risk on a continental scale.

This study aims to robustly predict how climate change will affect fascioliasis risk across Africa and Europe using a hybrid modelling framework that combines correlative and mechanistic approaches. We do so by developing an integrated risk index that predicts if environmental conditions are suitable for transmission of *F. hepatica* and *F. gigantica* based on three components: 1) the temperature dependence of liver fluke transmission [11,12,31], 2) the climatic suitability for the main intermediate host snail, and 3) the availability of aquatic habitats. Together, these represent different climatic and environmental requirements for completion of the liver fluke life cycle outside the main host. We then project this index under future climate scenarios to observe shifts in fascioliasis risk. By overlaying current and future risk maps with livestock density data layers, we highlight potential geographical hotspots where emerging fasciolosis risks could have the biggest impacts on livestock production. Lastly, to examine if climate change could increase the risk of hybridization between the two *Fasciola* species, we superimpose suitability maps for both species to identify overlapping areas where environmental conditions are favourable to transmission of both species.

## 2 Methods

We propose a risk index based on three components that capture the key environmental requirements for liver fluke transmission (Fig 1). The first component, termed “transmission capacity”, represents the temperature dependence of liver fluke transmission. It is estimated based on thermal response curves for important life history traits related to *F. gigantica* and *F. hepatica* transmission in the environment and the intermediate snail host. Secondly, to represent the requirements for presence of the suitable intermediate host snails, we use species distribution models (SDMs) to estimate climatic suitability for the most important intermediate host snails for each *Fasciola* species in Africa and Europe, namely *Radix natalensis* for *F. gigantica* and *Galba truncatula* for *F. hepatica* [32,33]. The final component reflects the requirement of the availability of stagnant water and is derived from hydrological model outputs. In the following sections, each of these components are described in detail.

**Fig 1.**
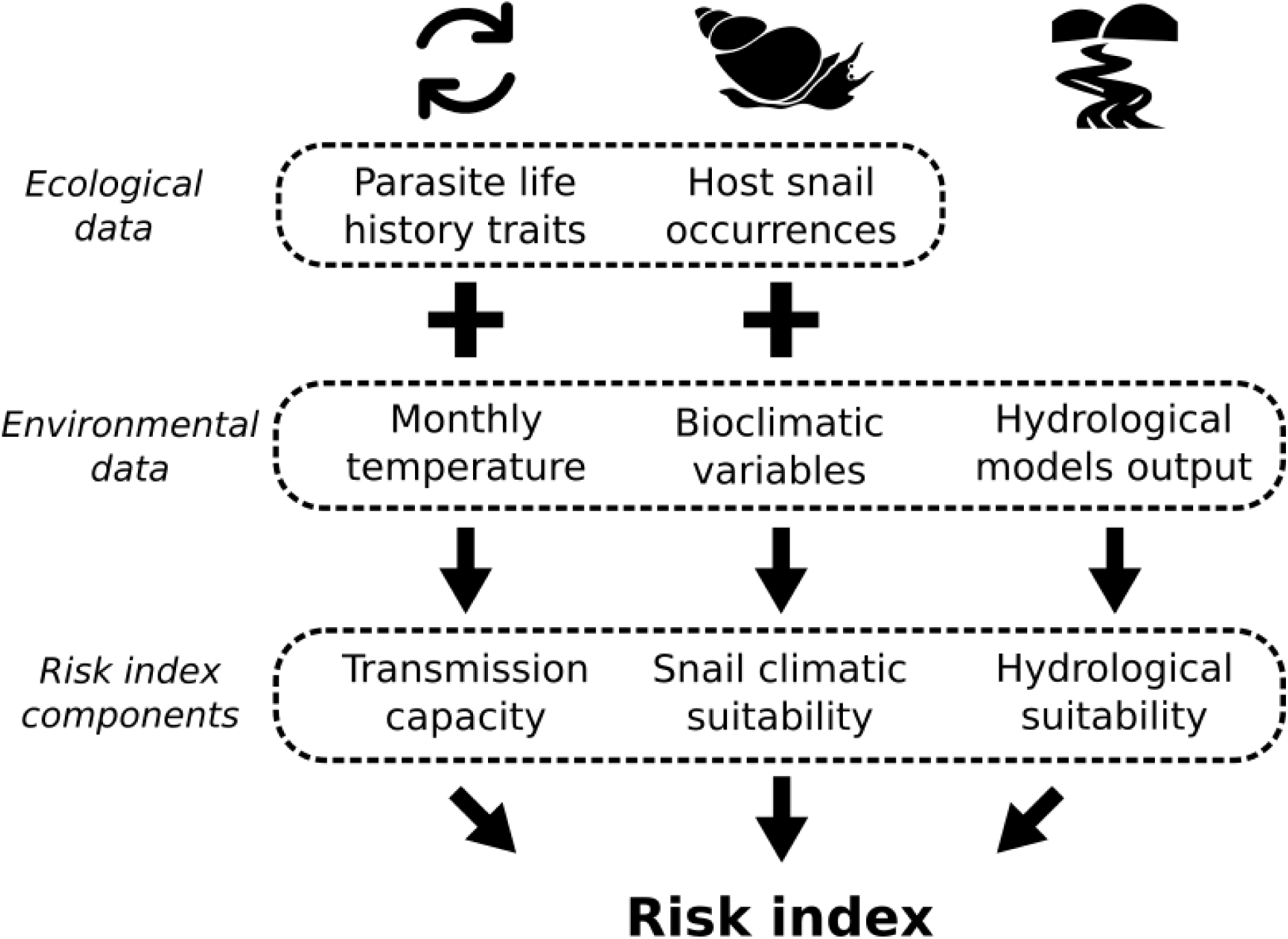
Components of the developed risk index for *Fasciola* transmission and the data used to generate them. Temperature-dependent transmission capacity is estimated based on data from laboratory-based experiments that measure temperature-dependent traits related to the transmission of *Fasciola* spp. These are then projected using monthly temperature data. Host snail climatic suitability is estimated from compiled occurrence data using bioclimatic variables as predictors. Finally, water availability is derived from the outputs of hydrological models.

### 2.1 Parasite life history trait data extraction and thermal response curve estimation

To estimate the temperature-dependence of transmission of *F. hepatica* and *F. gigantica,* we first estimated thermal response curves for traits related to the intramolluscan and free-living stages of their life cycle. To identify studies that include experimental data of relevant traits, we first obtained a list of references from recent systematic reviews by Dube et al. [12] and Modabbernia et al. [11]. We then identified further relevant articles from the reference lists of relevant papers (Table S1). Documents that were not available online were digitized from the collection of the Royal Danish Library.

We only included papers that reported on controlled temperature experiments, excluding reviews of earlier experiments, field experiments or observations, or laboratory experiments where temperature was not controlled. However, we did include studies that did not have thermal responses as their primary focus, even if they only used a single experimental temperature. If snails were subjected to other conditions than *Fasciola* infection, such as infection with another parasite, toxins, or desiccation, we used only data from the control group. Where data was only presented in figures, Plot Digitizer (https://sourceforge.net/projects/plotdigitizer/) was used to extract data points.

Using the extracted experimental data, we then fitted hierarchical Bayesian models for five *Fasciola* life history traits related to the environmental and intramolluscan stages of the parasite life cycle: egg hatching time; egg hatching success; infection efficiency (i.e. the proportion of miracidia successfully infecting a snail host); the prepatent period (i.e. the time from infection of the snail to onset of shedding); and cercarial release (i.e. the number of cercariae released from a single infected snail). We represented the hierarchical nature of the data, with repeated observations from each experiment, in hierarchical models, where the height of each curve was allowed to vary between experiments (see S1 Supplementary Materials for detailed descriptions of each model). The models were implemented in Julia v1.11 using the Turing library [34,35]. Models were sampled for 1000 iterations using the No U-Turn Sampling (NUTS) sampler [36] and we verified convergence using R-hat and effective sample size.

### 2.2 Estimation of the temperature dependence of parasite transmission

Next, we derived a measure we call “transmission capacity” from all five temperature-dependent traits. Transmission capacity is here defined as the relative number of cercariae expected to enter the environment after a single liver fluke egg is released, assuming the intermediate host is present. It thus measures how fast the parasite can proliferate at a given temperature. This measure is conceptually similar to “vectorial capacity”, a measure of transmission intensity of mosquito-borne diseases [37] and similar to earlier mechanistic models of fascioliasis transmission risk [29]. The temperature-dependent transmission capacity can be calculated by multiplying the estimated parasite egg survival, infection efficiency, survival inside the snail, and cercarial release:

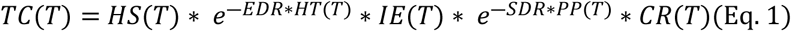

Where TC = transmission capacity; HS = hatching success, EDR = egg death rate, HT = hatching time, IE = infection efficiency, SDR = snail death rate, PP = prepatent period, CR = cercarial release, and T = temperature. Thus, all parameters are dependent on temperature, except for the egg and snail death rate, which were both fixed at 0.01 in our main model formulation (sensitivity analysis: see section 2 and Fig K in S1 Supplementary Materials).

As this model only accounts for the effects of temperature and assumes the presence of an aquatic habitat and suitable intermediate hosts, its output should be seen as a relative measure, rather than the actual number of cercariae to be expected in the environment. To reflect this, the transmission strength was normalized to between 0 and 1 for each sample in the posterior distribution. To project the transmission capacity, we calculated the monthly value using monthly temperature data and then averaged these over the entire year.

### 2.3 Snail intermediate host climatic suitability modelling

#### 2.3.1 Snail occurrence data

Occurrence records for the snail intermediate hosts *G. truncatula* and *R. natalensis* were extracted from three sources: GBIF; published literature on PUBMED; and finally previously unpublished records from the Freshwater Mollusc Collection of the Natural History Museum of Denmark. On GBIF, we performed a comprehensive search for lymnaeids recorded in Africa or Europe (see Data Availability for DOIs). GBIF records were cleaned by excluding records before 1980 to match the period for long-term climate averages and with coordinate uncertainty over 1000 metres. However, since very few records were available for Africa and the majority were from before 1980, we retained all African occurrence records. We additionally obtained previously unpublished occurrence records for lymnaeid snails in Africa from the Mandahl Barth Mollusc collection of the Natural History Museum of Denmark. Only species-level records were retained, and the same data cleaning procedure was applied as for GBIF data. Finally, we searched PUBMED for papers on *Fasciola* vector snails in Africa published between 2015 and 2025. Articles were then screened manually for relevance and snail species, and geographical location data were extracted. All data extraction procedures followed the Preferred Reporting Items for Systematic reviews and Meta-Analyses (PRISMA) guidelines. The full search query and inclusion protocol are given in S1 Supplementary Information (section 3.1).

After cleaning and spatial thinning, 3569 occurrences of *G. truncatula* and 672 occurrences of *R. natalensis* were used for fitting SDMs. Of the *G. truncatula* occurrences, only 49 were in Africa, while *R. natalensis* was only recorded in Africa (Fig L in S1 Supplementary Materials).

#### 2.3.2 Species distribution modelling

Species occurrence data is highly unevenly distributed in space due to uneven sampling efforts [38], especially in Africa [39]. Sampling background points (sometimes known as pseudo-absences) in a way that accounts for this bias is an effective way to mitigate the effects of sampling bias in models [40]. Therefore, we used a background sampling strategy where half of background points are sampled from occurrences of other lymnaeid species, and the other half is sampled randomly in space, to ensure areas unsuitable for all snail species such as deserts are represented. The total number of background points was double the number of occurrences. Furthermore, as the number of snail occurrences was highly unevenly distributed between Europe and Africa (Fig L in S1 Supplementary Materials), background points were sampled proportionally to the number of occurrence records on each continent (as *R. natalensis* only occurs in Africa, we only sample background points from Africa).

Environmental covariates were selected from the bioclimatic variables available on WorldClim [41]. We selected variables such that the correlation between all variables was low (all Pearson’s r < 0.7), and prioritized general variables (e.g. annual means) over monthly ones (e.g. precipitation of the wettest months), or variables that combine precipitation and temperature (e.g. mean temperature of the driest quarter) [42]. Using these criteria, we selected four bioclimatic variables as covariates: annual mean temperature (bio1), annual mean precipitation (bio12), temperature annual range (bio7), and the precipitation of the driest quarter (bio17). We fitted SDMs using Maxnet, which is a different implementation of the MaxEnt maximum entropy modelling software [43]. Maxnet was run with hinge features disabled and its regularization multiplier set to 3 to reduce overfitting, and default settings otherwise. To tests model fit, we performed 5-fold cross validation and measured the Area under the Receiver Operating Curve (AUC), and the True Skill Statistics (TSS) [44].

Cross-validated performance metrics showed that the SDMs captured the distributions of the vector snail species well (Table 1).

**Table 1.** Cross-validated performance of intermediate host species distribution models. Performance was measured using 5-fold cross-validation. Values shown are average performance on test data, with performance on training data in brackets. AUC = Area Under the receiver operator Curve. TSS = True Skill Statistic.

| Species | AUC | TSS |
| --- | --- | --- |
| *G. truncatula* | 0.76 (0.76) | 0.44 (0.43) |
| *R. natalensis* | 0.69 (0.70) | 0.33 (0.32) |

### 2.4 Hydrological suitability

The availability of standing water habitat is a requirement for the completion of the *Fasciola* life cycle. Hydrological discharge can serve as a proxy for the availability of aquatic habitats, as has been shown for mosquito larvae [45]. The hydrological models that estimate discharge take outputs from climate models, including gridded precipitation data, but also model hydrological flows such as evaporation, rivers, and irrigation. Here, we assumed that hydrological suitability in each grid cell follows the 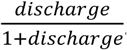 function. Hydrological suitability is thus 0 if no water flows through the grid cell, reaches 0.5 at a discharge of 1 mm/day and then asymptotically approaches 1. We set the 1 mm/day threshold following previously for modelling studies on aquatic mosquito habitats; however, hydrological suitability outputs are not particularly sensitive to the discharge threshold set [45,46]. Hydrological suitability was projected for each month using a monthly average of discharge (see section 2.6).

### 2.5 Combined Fascioliasis Risk index

Our novel transmission risk index for transmission of *F. hepatica* and *F. gigantica* is calculated from the three components described in the sections above:

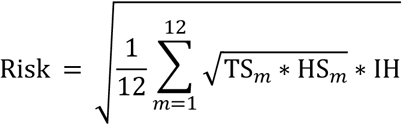

Where TS = temperature-dependent transmission capacity (section 2.3), HS = hydrological suitability (section 2.5), and IH = climatic suitability for the intermediate host (section 2.4), and *m* denotes each month. In other words, the risk index was computed by taking the geometric mean of transmission capacity and hydrological suitability for each month, calculating a mean value over the entire year, and then taking the geometric mean of this value and the climatic suitability for presence of the respective intermediate host. The risk index thus identifies areas where suitable temperatures for parasite transmission and freshwater availability simultaneously occur, and whose climate is suitable for the most intermediate host. Furthermore, since each component value is bound between 0 and 1, the risk index is also between 0 and 1.

### 2.6 Climatic and hydrological data

Gridded, bias-corrected historical climate data and global circulation model (GCM) output was downloaded from WorldClim [41] at 2.5 arc-minute resolution. Average temperature data was used for mapping parasite temperature suitability, and bioclimatic variables for modelling of habitat suitability of host snails. For forecasted data from both data sources, we downloaded data for the five GCMs that are available on ISIMIP: (GFDL-ESM4, IPSL-CM6A_LR, MPI-ESM1-2-HR, UKESM1-0-LL, and MRI-ESM2-0). Data was downloaded for two different Shared Socioeconomic Pathways (SSPs): SSP126, which assumes rapid action to bring down greenhouse gas emissions, and SSP370, which assumes medium-to-high emissions, and for two time periods: the middle of the century (2041-2060) and the end of the century (2081-2100). Long-term climatic averages for these time periods are available directly from WorldClim.

Discharge data from hydrological model output was downloaded from ISIMIP at 0.5° resolution [47]. We downloaded monthly model output for all four Global Hydrological Models (GHMs) available (CWatM [48], JULES-W2 [49,50], MIROC-INTEG-LAND [51], and WaterGAP2-2e [52]). For future data, se downloaded outputs for the same four GHMs, which were forced by the same five GCMs as for climate data. We generated monthly averages by averaging monthly outputs over the same years as for climate data (1971-2000, 2041-2060, and 2081-2100) for each month. Discharge data was converted from m^3^/s to mm/day by dividing by the cell area of each grid cell. For calculating the final risk index, we then disaggregated to match the 2.5 arc-minute resolution of the climate data.

Some of the latest generation of climate models overestimate the amount of warming resulting from greenhouse gas emissions, and experts advise to act with care when including these models [53,54]. UKESM1-0-LL has a transient climate response of 2.77°C, which is outside of the “very likely” range as assessed by the IPCC [53]. We therefore excluded this model from the average forecasts.

### 2.7 Livestock data and plotting overlap

In order to identify hotspots of overlap between high liver fluke risk and high livestock densities i.e. where the impact on livestock production might be highest, we superimposed the risk index surfaces for each *Fasciola* species with sheep and cattle densities. Likewise, we superimposed modelled *F. hepatica* and *F. gigantica* risk surfaces under current and future conditions to identify areas of overlapping distributions between the two species, to investigate where hybridization could potentially take place now and in the future. Downscaled, gridded datasets for sheep and cattle densities for 2020 were downloaded from the GLW4 dataset [55]. These data are based on census data and then downscaled to 5 arc-minutes [56].

## 3 Results

### 3.1 Thermal biology of *Fasciola* transmission

To investigate the role of temperature in transmission of *F. hepatica* and *F. gigantica,* we estimated thermal performance curves for five parasite life history traits and used a life cycle model to estimate the temperature dependence of the entire transmission cycle (Eq. 1). The thermal response curves derived from these data points show that *F. hepatica* life history traits peak at lower temperature than *F. gigantica* traits, except for infection efficiency, which peaks at similar temperatures (Fig 2, Section 1 and Figs A-J in S1 Supplementary Materials). The parasite egg hatching time increases rapidly around 10°C for *F. hepatica* and around 15°C for *F. gigantica* (Fig 2A). Cercarial release for *F. hepatica* peaks around 18°C and is negligible above approximately 25°C, limiting transmission strength above these temperatures (Fig 2D). For *F. gigantica*, infection efficiency, hatching success, and cercarial release all decline strongly around 35°C. All data points and prior and posterior distributions for each parameter are provided in the Figures S1-S10 and Tables S2-S6.

**Fig 2.**
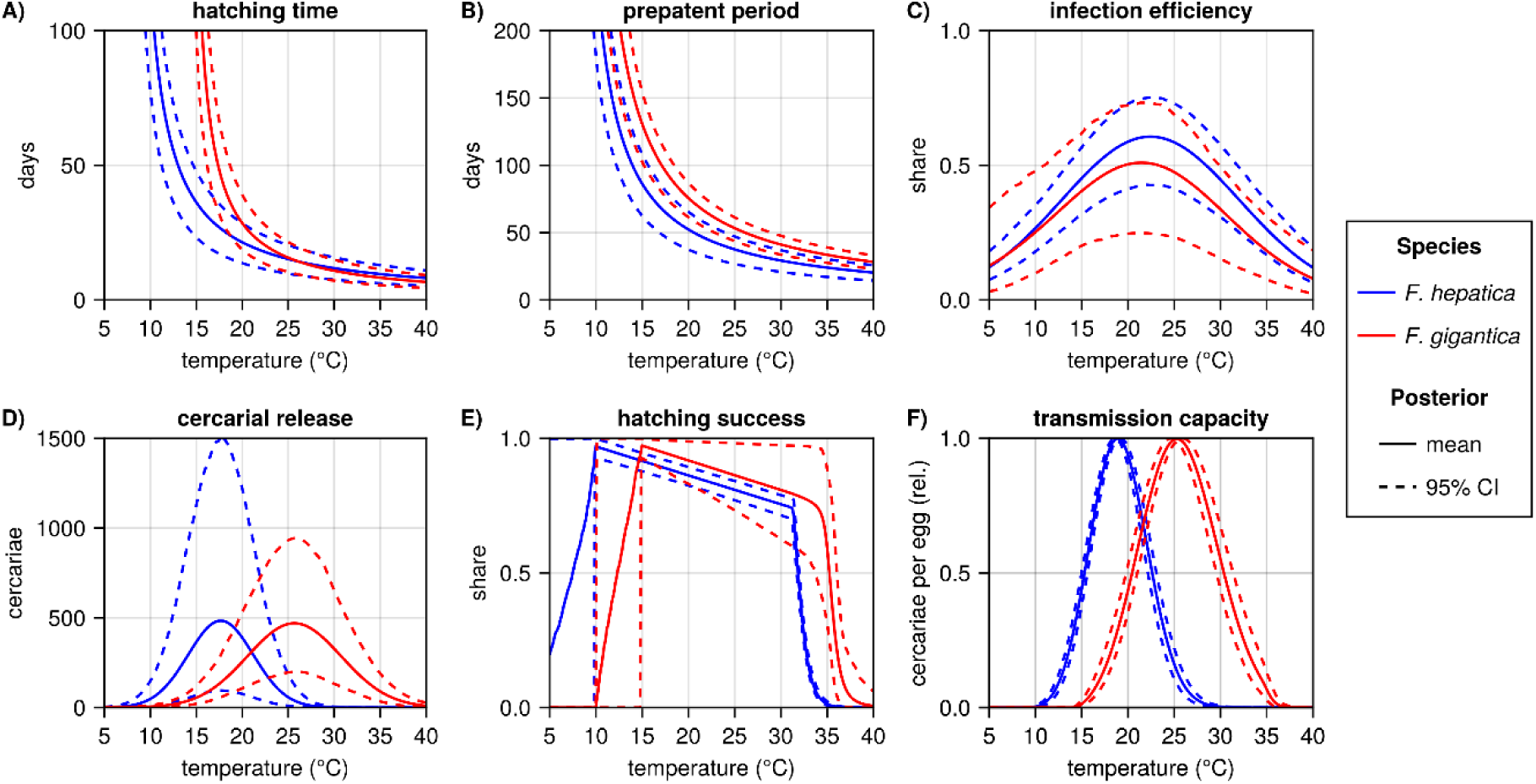
Posterior estimates for thermal performance curves for five life history traits and estimated transmission strength for *Fasciola hepatica* (blue) and *Fasciola gigantica* (red). Mean estimates (solid lines) are plotted along with and 95% credible intervals (dashed lines). The five life history traits are A) time to parasite egg hatching, B) prepatent period, i.e. time from snail infection to the onset of cercarial shedding, C) the probability of successful snail infection by a single miracidium, D) the number of cercariae released by an infected snail, E) the share of parasite eggs that successfully hatch. F) the temperature-dependent transmission strength, as estimated by the transmission cycle model based on the five life history traits (main text Eq. 1), shows that *F. hepatica* is adapted to colder temperatures than *F. gigantica*.

These life history traits were then used to parameterise a life cycle model (Eq. 1) that estimates how transmission strength depends on temperature. The resulting estimates of temperature-dependent transmission strength confirm that the optimal temperature for *F. hepatica* transmission is much lower than that of *F. gigantica* transmission, at 18.9°C (95% CI 18.6-19.2) and 25.3°C (95% CI 24.7-25.9), respectively (Fig 2F). Similarly, *F. hepatica* transmission is possible between around 10°C to around 30°C, whereas *F. gigantica* transmission is possible between around 15°C to around 36°C.

### 3.2 *Fasciola* transmission risk

We combined spatial estimates for the climatic suitability for the main snail intermediate host, temperature-dependent transmission strength, and availability of water into a novel index for transmission risk for *F. hepatica* and *F. gigantica* and projected this for Europe and Africa under current and future climatic conditions. The resulting projections show that the transmission risk for *F. hepatica* is substantial in most of Europe under current environmental conditions, driven by high climatic suitability for the snail intermediate host *G. truncatula* (Fig 3A-D). Climatic suitability for *G. truncatula* is particularly high in western Europe, including the British Isles and Iceland, while it is low in southern Spain, limiting transmission risk there (Fig 3C). Temperature-dependent transmission strength in Europe is relatively low, which may be due to the seasonal transmission pattern, with cessation of transmission in winter months. In Iceland and northern Scandinavia, temperatures are completely outside of the thermal limits of transmission, which is why the risk index shows there is no transmission risk there. In Africa, *F. hepatica* hotspots are found in the East-African highlands and in Southern Africa. *F. hepatica* transmission risk is limited in most of Africa because of the low climatic suitability for *G. truncatula*, while temperatures are suitable for transmission in most of Africa except for the Sahel.

**Fig 3.**
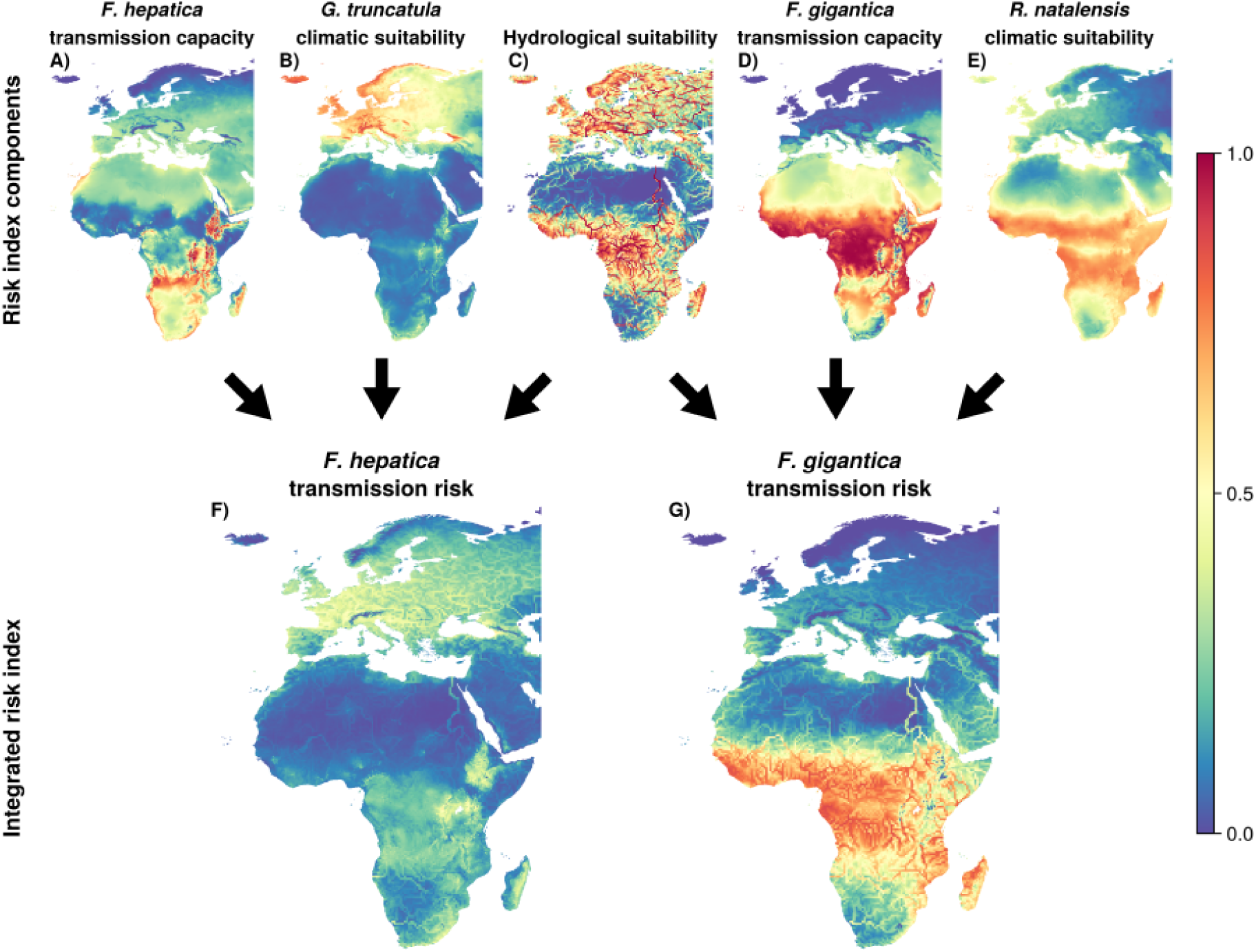
Projected transmission risk for F) *Fasciola hepatica* and G) *Fasciola gigantica* and the risk index components (A-E) under current climate conditions. **A)** Temperature-dependent transmission capacity for *F. hepatica*. **B)** Climatic suitability for *Galba truncatula*, the most important intermediate host for *F. hepatica*. **C)** Hydrological suitability, which represents availability of aquatic habitats, is used in the risk indices for both *Fasciola* species. **D)** Temperature-dependent transmission capacity for *F. gigantica*. **E)** Climatic suitability for *Radix natalensis*, the most important intermediate host for *F. gigantica*. **F)** Transmission risk index for *F. hepatica*, which shows transmission risk is high across Europe and in the East-African highlands. **G)** The transmission risk index for *F. gigantica* is high in tropical Africa and very low elsewhere.

For *F. gigantica*, the risk index shows that transmission risk is high in most of Africa, except for desert areas and the Mediterranean coast (Fig 3G). In desert areas, both climatic suitability for the intermediate host snail *R. natalensis* and hydrological suitability are very low (Fig 3A and Fig 3F). In the tropical regions around the equator, including in West, Central, and East Africa, temperatures are close to optimal for transmission of *F. gigantica* (Fig 3E). We predict very low risk of *F. gigantica* transmission across Europe, driven by both low temperature-dependent transmission strength and low climatic suitability for *R. natalensis*. However, temperatures are high enough for *F. gigantica* transmission in some parts of Southern Europe, indicating transmission could be possible if an intermediate host were also present.

### 3.3 Fascioliasis transmission risk under climate change

We then projected the transmission risk index under climatic conditions at the middle and end of the century under the low-emission SSP126 scenario and the SSP370 scenario with medium-to-high emissions. Predictions under the SSP370 scenario show the risk of *F. hepatica* transmission is projected to decline drastically across the African continent over the course of this century, with only a few pockets of risk remaining in the highlands and in South-Africa (Fig 4). Risk is also projected to decrease in Southern Europe, but to increase slightly in Northern Europe, especially in colder areas such as the British Isles, Scandinavia, and Iceland, which historically has been free of *Fasciola* [57]. Under the SSP126 scenario, which assumes drastic cuts to emissions, the direction of the changes is similar, but they are less pronounced, and risk is projected to stabilise after the middle of the century (Fig M in S1 Supplementary Materials).

**Fig. 4.**
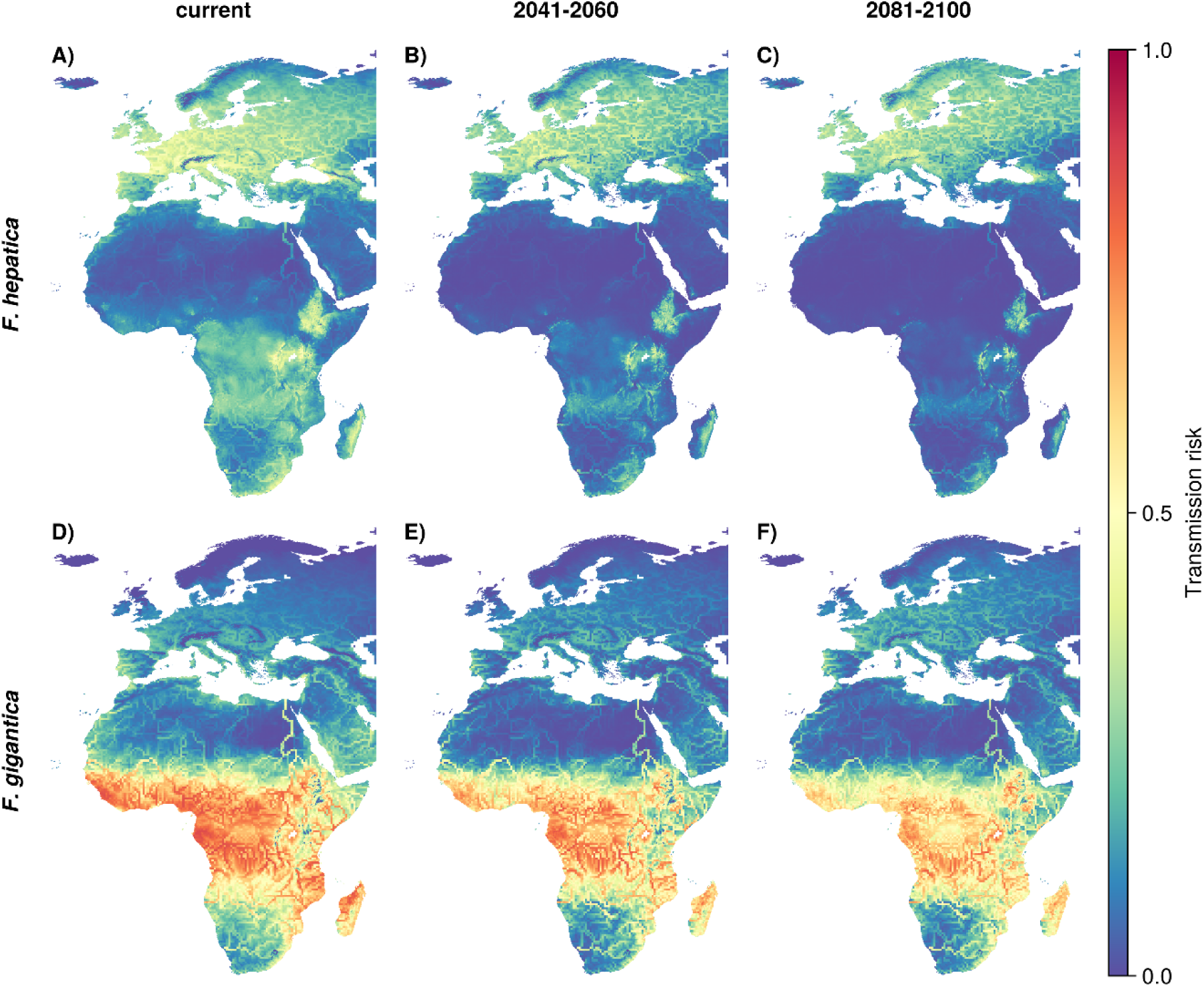
Current and future transmission risk for *Fasciola hepatica* and *Fasciola gigantica*, as estimated by the transmission risk index. Projection assume the SSP370 scenario with high emissions. **A)** Current transmission risk for *F. hepatica*, as shown in Fig 3F. **B)** Projected *F. hepatica* transmission risk for the middle of the century (2041-2060). C) *F. hepatica* transmission risk projected for the end of the century 2081-2100. Risk is projected to decrease across most of the range of *F. hepatica*, especially in Africa. **D)** Current transmission risk for *F. gigantica*, as in shown in Fig 3G. **E)** Projected *F. gigantica* transmission risk for 2041-2060. F)*F. gigantica* risk for 2081-2100. The transmission risk is projected to decrease under future climatic conditions.

For *F. gigantica*, projections suggest that transmission risk will decline in West and Central Africa and increase in parts of East Africa. However, most of Africa south of the Sahara will continue to be at risk of *F. gigantica* transmission by the end of the century, even under the SSP370 scenario. The risk of *F. gigantica* transmission in Southern Europe is projected to remain low.

The changes in projected risk are driven by divergent changes in transmission strength and snail host suitability. Temperature-dependent transmission strength for *F. hepatica* is projected to increase in Europe and decrease in most of Africa (Figs N-O in S1 Supplementary Materials). However, this is offset by decreasing environmental suitability for its vector snail *G. truncatula* across most of its range (Figs P-Q, top row in S1 Supplementary Materials). For *F. gigantica,* sharply decreasing transmission strength drives the lower transmission risk in West-Africa and the Sahel, indicating temperatures in these areas will become less suitable for transmission of *F. gigantica*. Transmission risk is also forecast to stay stable in most of East Africa, increase in the East-African highlands and Europe. Our forecasts indicate little change in the environmental suitability for *R. natalensis*, the main intermediate host snail for *F. gigantica* (Figs P-Q, bottom row in S1 Supplementary Materials).

### 3.4 Impacts on hybridization risk and overlap with livestock

The emergence of hybrids or intermediate forms of *F. hepatica* and *F. gigantica* is a risk to disease control, as such forms can have advantageous traits and increased zoonotic potential [58]. To map areas where both *Fasciola* species can be transmitted, and where the risk for hybridization is therefore high, we superimposed the transmission risk indices of the two *Fasciola* species under current and future climatic conditions (Fig 5). Currently, in Southern Africa, Central Africa, and the East-African highlands are all areas where the risk index projections show that high risk for transmission with both *F. hepatica* and *F. gigantica*. However, under future climatic conditions, the area of overlap will shrink significantly. This is mainly driven by the declining transmission risk of *F. hepatica*. No new areas of overlap are predicted to emerge in Europe or Africa under the investigated climate change scenarios.

**Fig 5.**
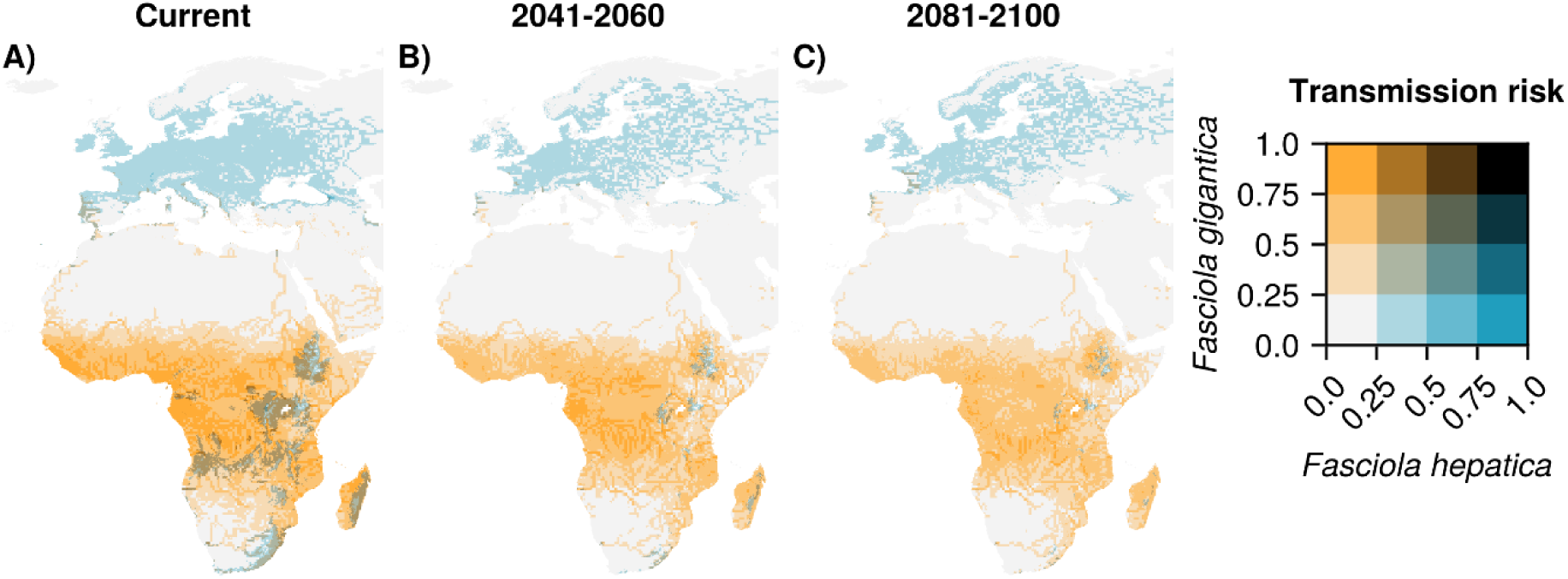
Current and projected future overlap of transmission risk of *F. hepatica* and *F. gigantica*. Transmission risk (as presented by the developed risk index) for *F. gigantica* is shown in orange and for *F. hepatica* in blue, where dark colours identify area with overlapping risk (i.e. risk index at least 0.25 for both species). **A)** Current areas where transmission for these species is both high include areas in Southern Africa and the East-African highlands. **B-C)** Under the SSP370 scenario with high emissions, the area of overlap is expected to decrease drastically in the future.

Given the big impacts of fascioliasis on livestock farming and the role of livestock as a zoonotic reservoir, we then identified areas with both a high risk for *Fasciola* transmission and high cattle and sheep densities. Currently, high risk of *F. gigantica* transmission overlaps with high density of both cattle and sheep in the Sahel region, East-Africa, and the Great Lakes region (Fig 6). Projected *F. hepatica* risk overlaps with high densities of sheep in the British Isles, parts of Mediterranean Africa, and Southern Africa, as well as cattle in Western Europe and the East-African highlands (Fig 6).

**Fig 6.**
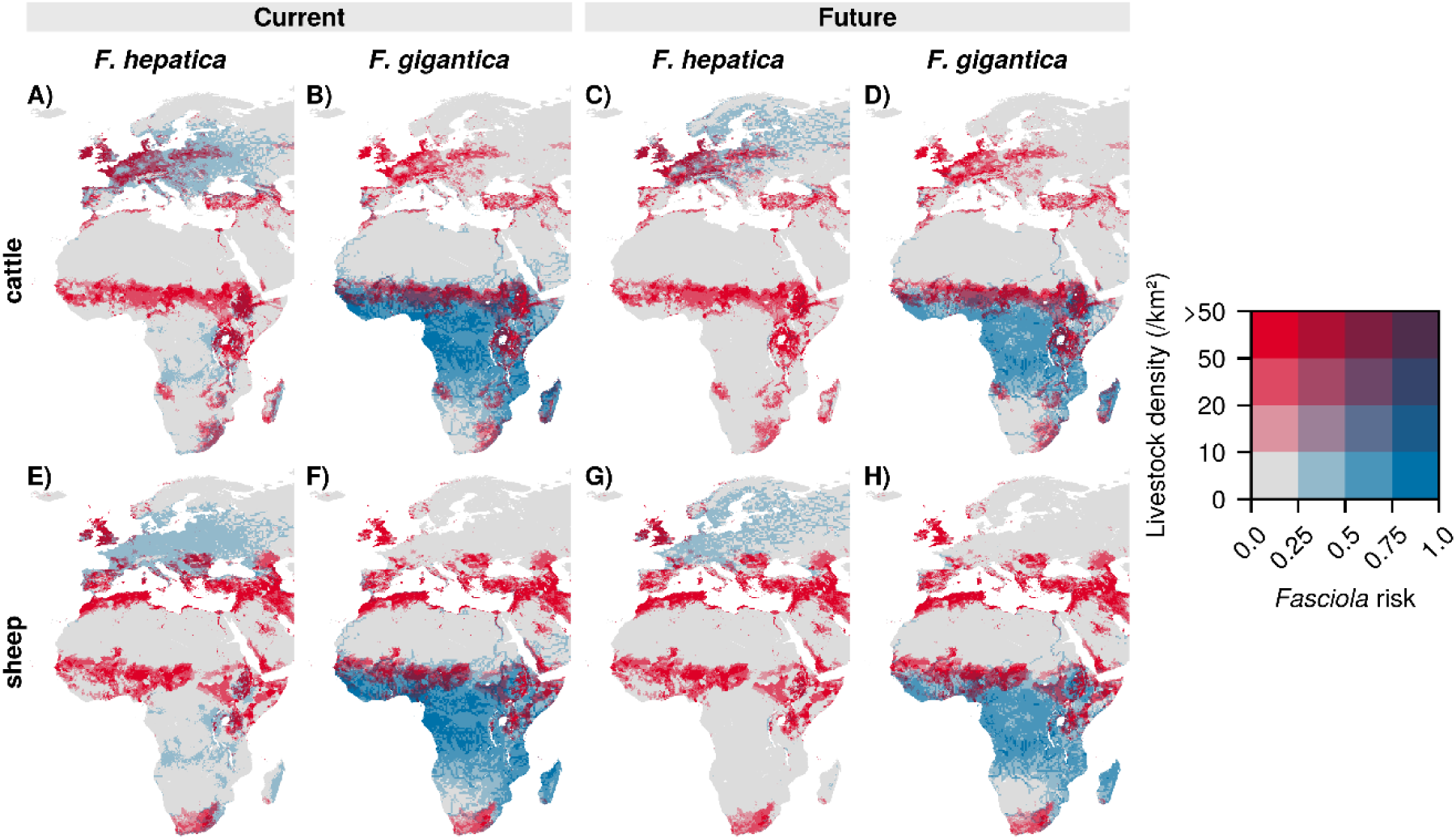
Current and future overlap of risk of *Fasciola* transmission and sheep and cattle densities. Livestock densities (in number of animals per square km) are shown in red and transmission risk in blue shades. The top row (A-D) shows cattle densities, and the bottom row (E-H) sheep densities. Current transmission risk is shown on the left half of the figure (*F. hepatica* A and E, *F. gigantica*, B and F). Forecasted future risk (under the SSP370 scenario) is shown in the right half of the figure (*F. hepatica* C and G, *F. gigantica* D and H).

Under climate change, *F. hepatica* transmission risk will remain high or even increase in areas of Western Europe where cattle and sheep densities are high, including on the British Isles. However, in Southern Europe, including Spain, risk is decreasing in areas with a high density of sheep. For *F. gigantica*, decreasing suitability is especially notable in the West-Africa and the Sahel region. However, an important caveat to these projections is that we have not modelled how climate change could affect livestock densities including the influence of secondary animal hosts, such as horses, mules, camels, and wild ruminants at local levels [59].

## 4 Discussion

In this study, we investigated the possible effects of climate change on *Fasciola* transmission across Africa and Europe. Our findings indicate decreasing risk of *Fasciola* transmission under climate change across most of the range of both *F. hepatica* and *F. gigantica*. However, in northern Europe, the risk of *F. hepatica* transmission might increase, with new areas at risk emerging in Scandinavia and Iceland. The shifting risk of fascioliasis could have significant impacts in northern Europe, where there are both high densities of sheep and cattle and increasing transmission risk, as well as the Sahel region, where sheep densities are high, and risk is decreasing. Finally, the area of overlapping transmission risk of *F. gigantica* and *F. hepatica*, where hybridization might occur, is projected to decrease because of climate change.

To make these predictions, we developed a new index of *Fasciola* transmission risk by combining an estimate of temperature-dependent transmission strength derived from life history traits and a life cycle model, water availability form hydrological models, and climatic suitability for the most important snail intermediate hosts. Combined, the different components of the index represent the whole liver fluke live cycle, except for the stages within the vertebrate host, which are less sensitive to changes in the environment. It also integrates multiple modelling approaches and data sources, arguably making it more robust, while also being applicable on large spatial scales due to its generality and relative simplicity [60,61]. This approach is a major advance over earlier studies on the changing risk of *Fasciola* transmission under climate change, which have either been confined to local scale, or have used much simpler models such as the Ollerenshaw index [23].

In addition, we provide novel estimates for the thermal performance curves for five life history traits of *F. hepatica* and *F. gigantica* based on experimental data. From these, we derived estimates for how transmission depends on temperature. Our estimate that *F. hepatica* transmission is possible from around 10°C and peaks at around 18°C is close to earlier estimates [23,62]. For *F. gigantica,* we estimate transmission is possible from around 15°C, which is also similar to what has been used in earlier models [18,21]. However, these studies did not estimate an optimal and maximum temperature for *F. gigantica* transmission, which we put at around 25°C and 35°C, respectively. These updated estimates can be used to parameterise other fascioliasis risk models, including local forecasting systems.

The fascioliasis risk we predict under current climatic conditions generally aligns well with the known patterns of fascioliasis transmission [63]. In Africa, *F. gigantica* is known to be widespread while *F. hepatica* is restricted to more temperate areas and highlands, which is consistent with our risk index [64–66]. In Europe, our projection that Western Europe is particularly at risk of *F. hepatica* also aligns with earlier correlative modelling [14]. However, high levels of fascioliasis have also been observed in some areas where we project low risk, such as Scotland [67]. This shows that while our risk index is consistent with continental-scale dynamics, its ability to predict on the local scale might be limited.

Our finding that fascioliasis risk will decrease because of climate change stands in contrast with earlier studies on the risk of *F. hepatica* transmission under climate change scenarios in Europe, which have predicted risk to increase across the continent by applying the Ollerenshaw index to future climate projections [23,24]. This difference is likely due to the different modelling approach we use. While we do project increasingly suitable temperatures for transmission, we also predict decreasing suitability of intermediate snail hosts, which these earlier papers do not account for. In Africa, earlier studies that predict the climate change impacts on *Fasciola* transmission have only investigated changes at the local or national scale [18,21,68]. To the best of our knowledge, this study is the first to estimate *Fasciola* transmission risk across Africa, showing decreasing risk because temperatures will be less suitable for transmission with climate change.

By overlapping the modelled risk index surfaces for both *F. hepatica* and *F. gigantica*, we also investigated how climate change might affect the co-occurrence of these species and thus the potential for hybridization to occur. Our results suggest that the overlap in these distributions will decline with climate change, diminishing the risk of hybridization. However, earlier hybridization and introgression events have been linked to livestock movement, which is steadily increasing [58,69]. Thus, hybridization will remain an important risk despite climate-driven range shifts, especially if global livestock movement keeps increasing. More research is needed on intermediate forms or hybrids, as their adaptive capacity and fertility is still unclear [58,70].

A limitation of our study is that the risk index we have developed focuses only on the environmental parts of the *Fasciola* life cycle and therefore may not be directly translatable to actual prevalence. Whether a higher risk index will result in a higher prevalence of fascioliasis might depend on a multitude of additional factors. Locally, at smaller spatial scales, non-climatic factors such as grazing practices, local topography, or drainage often found to be more important predictors for fascioliasis prevalence in livestock [71–75]. Both within and between Africa and Europe, such local factors are widely different, resulting in different prevalences regardless of climatic factors. Similarly, the risk of human fascioliasis is closely linked to the consumption of specific freshwater plants such as watercress or unrinsed vegetables, which is much more common in some countries than others [59,76]. Our approach is therefore mostly useful to understand continental-scale dynamics under global change, but cannot replace systems to predict seasonal risk on a smaller, local scales [29]. Conversely, as climate change is pushing temperatures into unknown territories [77], it is important to ensure that local risk assessments or forecasting systems are based on as complete ecological information as possible, and are thus able to forecast beyond previously observed conditions. The modelling approaches and thermal performance curves we present here could therefore be incorporated into local prediction systems.

Furthermore, we assume that both *Fasciola* species and their intermediate hosts have the same environmental tolerances throughout the study region (i.e. niche conservatism [78]). While niche conservatism is a foundational assumption in SDMs [78,79], genetic adaptations to local climatic conditions may be important and have been shown in freshwater snails [80]. Our recent work suggests such adaptations may also be present in *R. natalensis* and *Radix balthica* across Europe and Africa [81]. For *G. truncatula* specifically, recent work has suggested that this amphibious snail is made up of a complex of cryptic species and that snails identified as *G. truncatula* in Africa may be distinct from *G. truncatula* in Europe [82]. In addition, liver flukes have different transmission strategies in different climates; in Europe, the overwintering of eggs may be important for spring infection of *F. hepatica* [83,84], while aestivating infected snails are key to maintaining transmission cycles in many settings in Africa [85,86]. More work is needed to investigate how such variability might help *Fasciola* parasites and their intermediate hosts adapt to climate change in the future.

*Fasciola* parasites can use different intermediate host species, which could imply a different climate change response [87]. One of these is the invasive and highly resilient *Pseudosuccinea columella*, which is an intermediate host for both *F. hepatica* and *F. gigantica* and a serious concern for fascioliasis risk [88,89]. Even though its competence as a *F. hepatica* intermediate host might be lower than that of *G. truncatula* [90], recent evidence suggests it might be playing a key role in *F. gigantica* transmission in South-Africa [91]. The spread of *P. columella* might reduce *Fasciola* transmission if it displaces more competent local vectors [92], or it might maintain transmission if it adapts to changing climatic conditions more easily. In the Americas and in Asia, *P. columella* and other lymnaeid species act as the intermediate hosts for *Fasciola* species [87]. Thus, future research is needed to elucidate how climate change impacts interact with inter- and intraspecific variation of host snails and the spread of invasive snails. Such studies could draw on the modelling approach and thermal performance curves presented here while accounting for the biology of the local intermediate host snails.

Finally, while the projected risk index corresponds well to known occurrence patterns, we have not formally validated our risk index. Such validation would be challenging for a number of reasons, including the big differences in fascioliasis prevalence between farms in a single region [14], and the uncertainty about where infections are acquired for studies that register fascioliasis prevalence at abattoirs (e.g. [93]). There are also big discrepancies in data availability between Europe and Africa. In Europe, the international research projects with standardized procedures across countries have done much to elucidate the spatial patterns of *F. hepatica* prevalence [94]. In Africa, however, research has been smaller in scale, and thus fascioliasis prevalence is much more uncertain [64,95]. Leveraging the available prevalence data to map fascioliasis prevalence throughout Africa is a research avenue that is yet to be explored and could be a valuable addition to this study.

## 5 Conclusion

In this study, we developed a novel risk index for *F. hepatica* and *F. gigantica* transmission to investigate the potential impacts of climate change on the fascioliasis risk across Africa and Europe. This index is based on both the thermal biology of the *Fasciola* parasites and the environmental suitability of their most important host snails. We highlight uneven impacts of climate change on fascioliasis risk, with increasing risk for *F. hepatica* at higher latitudes and decreasing risk for both *F. gigantica* and *F. hepatica* in most other areas. In Northern Europe, where sheep and cattle densities are high, increasing transmission risk could have major impacts. Our study offers valuable insights into continental-scale dynamics under global change, but we recognise that local factors also play a key role in determining fascioliasis risk. By developing a new hybrid modelling framework and providing insights into the thermal biology of the *Fasciola* lifecycle, this work lays the groundwork for the development of more robust local risk indices and early warning systems.

## 6 Data and code availability

All code used and instructions to reproduce all analysis performed are available on GitHub at https://github.com/tiemvanderdeure/FasciolaDistribution. Hydrological model outputs are available on ISIMIP (via https://files.isimip.org/). Climate data is available on WorldClim at www.worldclim.org. Occurrence data from GBIF can be permanently accessed at https://doi.org/10.15468/dl.qekchb and https://doi.org/10.15468/dl.jqdg8p. Snail occurrence data extracted from the literature and the Natural History Museum of Denmark, as well as experimental life history traits, are included in the GitHub repository. Gridded livestock data is available from the FAO (https://data.apps.fao.org/catalog//iso/9d1e149b-d63f-4213-978b-317a8eb42d02).

## Acknowledgements

We gratefully acknowledge the researchers whose fieldwork and experimental studies provided the foundation for the data presented in this work.

This project has received funding from the European Union’s Horizon 2020 research and innovation program under grant agreement No. 101000365 (PREPARE4VBD).

## Author contributions

TvdD, ASS, MES, MC, SK, and SM conceptualized the study and designed the methodology. ASS, MES, SK, MC and SM acquired funding and administrated the project. ASS and DNB supervised the work. TvdD, RM, JW, BON, MS, AD, and TM contributed to data compilation and curation. TvdD carried out the analysis, visualized the results, and wrote the initial draft. ASS, SM, DNB, and TM critically reviewed the initial draft. All authors have read and approved the final draft of the manuscript.

## 10 Supplementary materials captions

**S1 Supplementary Materials contains:**

**Figures A-S**

**Fig A. Data sources and prior and posterior distributions for F. gigantica hatching time.** Upwards arrows indicate a truncated value, where the experiment was stopped before hatching took place.

**Fig B. Data sources and prior and posterior distributions for *F. hepatica* hatching time.**

**Fig C. Data sources and prior and posterior distributions for *F. gigantica* prepatent period.**

**Fig D. Data sources and prior and posterior distributions for F. hepatica prepatent period.**

**Fig E. Data sources and prior and posterior distributions for *F. gigantica* infection efficiency.**

**Fig F. Data sources and prior and posterior distributions for *F. hepatica* infection efficiency.**

**Fig G. Data sources and prior and posterior distributions for *F. gigantica* cercarial release.**

**Fig H. Data sources and prior and posterior distributions for *F. hepatica* cercarial release. Fig I. Data sources and prior and posterior distributions for *F. gigantica* hatching success Fig J. Data sources and prior and posterior distributions for *F. hepatica* hatching success**

**Fig K**. **Sensitivity analysis for the effect of different snail and egg death rates on the transmission strength index.**

**Fig L. Occurrence records of *Radix natalensis*, *Galba truncatula*, and other lymnaeid snails.** Other lymnaeid occurrences were used as background points for species distribution modelling.

**Fig M. Projected *Fasciola* transmission risk under the SSP126 scenario with low emissions.**

**Fig N. Projected temperature-dependent transmission potential under the SSP126 scenario with low emissions.**

**Fig O. Projected temperature-dependent transmission potential under the SSP370 scenario with high emissions.**

**Fig P. Projected snail host suitability under the SSP126 scenario with low emissions. Fig Q. Projected snail host suitability under the SSP370 scenario with high emissions.**

**Fig R. Projected transmission risk for F. hepatica under each combination of global hydrological model (rows) and global circulation model (columns).** UKESM1-0-LL is known to predict warming beyond the plausible range and was therefore excluded from the mean estimates presented in the paper.

**Fig S. Projected transmission risk for F. gigantica under each combination of global hydrological model (rows) and global circulation model (columns).**

**Tables A-F**

**Table A. Studies with experimental life history data**

**Table B. Prior and posterior distributions for hatching time**

**Table C. Prior and posterior distributions for prepatent period**

**Table D. Prior and posterior distributions for infection efficiency**

**Table E. Prior and posterior distributions for cercarial release**

**Table F. Prior and posterior distributions for hatching success**

## Notes

### Competing Interest Statement

The authors have declared no competing interest.

### Summary of Updates

Supplementary materials were added. The main manuscript file is unchanged.

