## Supplementary material for "Novel integrated risk index reveals uneven impact of climate change on fascioliasis risk across Africa and Europe": S1 Supplementary Materials

### 1 Fasciola life history traits

Life history traits relating to the life cycles of *Fasciola gigantica* and *Fasciola hepatica* were extracted from sources during a thorough literature search. Table S1 shows an overview of sources that were used.

**Table A: Studies with experimental life history data**

|  | | |
| --- | --- | --- |
| **Citation** | **Species** | **Trait** |
| Abrous et al. (1999) | hepatica | prepatent_period, infection_efficiency, cercarial_release |
| Al-Habbib & Grainger (1983) | hepatica | prepatent_period, cercarial_release |
| Al-Habbib & Al-Zako (1981) | hepatica & gigantica | hatching_success, prepatent_period |
| Al-jibouri et al. (2010) | gigantica | hatching_time, hatching_success |
| Al-Jibouri et al. (2011) | gigantica | hatching_time, prepatent_period |
| Alicata (1938) | gigantica | hatching_time |
| Bitakaramire (1968) | gigantica | hatching_time, prepatent_period, cercarial_release |
| Christensen et al. (1976) | hepatica | infection_efficiency |
| Claxton et al. (1999) | hepatica | hatching_time |
| Connolly (1979) | hepatica | hatching_time |
| Dar et al. (2004) | gigantica | prepatent_period, cercarial_release |
| Dinnik & Dinnik (1963) | gigantica | hatching_time, prepatent_period |
| Dinnik & Diknik (1964) | gigantica | infection_efficiency |
| Dreyfuss et al. (1999) | hepatica | prepatent_period, infection_efficiency |
| Gold & Goldberg (1979) | hepatica | infection_efficiency |
| Grétillat (1961) | gigantica | hatching_time |
| Guobadia et al. (1996) | gigantica | prepatent_period, cercarial_release |
| Güralp et al. (1964) | gigantica | hatching_time |
| Kendall & McCullough (1951) | hepatica | cercarial_release |
| Moazeni et al. (2010) | hepatica | hatching_success |
| Nice (1979) | hepatica | prepatent_period, cercarial_release |
| Nice & Wilson (1974) | hepatica | hatching_time |
| Over (1982) | hepatica & gigantica | hatching_time, prepatent_period |
| Phalee et al. (2015) | gigantica | hatching_time |
| Raman et al. (2012) | gigantica | hatching_success |
| Rao (1966) | gigantica | hatching_time |
| Roberts (1950) | hepatica | hatching_time, prepatent_period, cercarial_release, hatching_success |
| Rondelaud et al. (2013) | hepatica | prepatent_period, infection_efficiency, cercarial_release |
| Ross & McKay (1929) | hepatica | hatching_time |
| Rowcliffe & Ollerenshaw (1960) | hepatica | hatching_time |
| Shalaby et al. (2004) | gigantica | prepatent_period, cercarial_release |
| Sharma et al. (1989) | gigantica | hatching_time |
| Smith (2016) | hepatica | hatching_time, hatching_success |
| Soliman (2009) | gigantica | infection_efficiency, cercarial_release |
| Tagle (1944) | hepatica | hatching_time |
| Wilson & Taylor (1978) | hepatica | infection_efficiency |

Using this data, we fit Bayesian models to the data using selected equations. To limit the amount of bias from choice of equation, we used the same equation for both species for each trait. We also used identical priors. In selecting the equation, we favoured simplicity and biological realism. This section will present the data used, model formulation, and prior and posterior distributions of variables for each life history trait. Models were fit in Julia v1.11 using the Turing probabilistic inference library.

#### 1.1 Hatching time

For hatching time, we use a growing degree day model, defined as:

$$HT\left( T \right)= \left\{ \begin{aligned} \frac{days}{T-Tmin}, &T>Tmin \\ 0, &T \leq Tmin \end{aligned} \right.$$

Where HT is the time to hatching, *days* is the number of growing degree days, and *Tmin* is the minimal temperature required for development.

We included 11 studies for *F. gigantica* and 9 studies for *F. hepatica* that recorded this trait*.*

Some studies were stopped before hatching took place. In this case, we used a truncated distribution.

In the Bayesian model, *days* was a random effect, allowed to vary between studies. *Tmin* was assumed to be equal between all the studies.

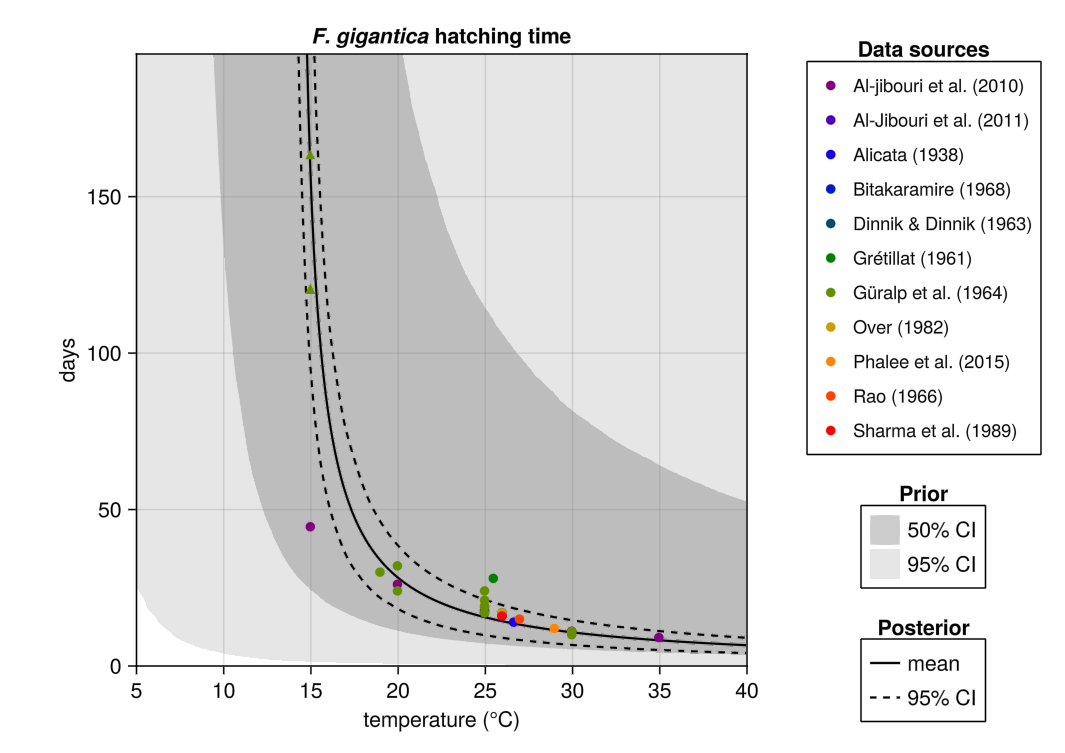

**Fig A. Data sources and prior and posterior distributions for *F. gigantica* hatching time**. Upwards arrows indicate a truncated value, where the experiment was stopped before hatching took place.

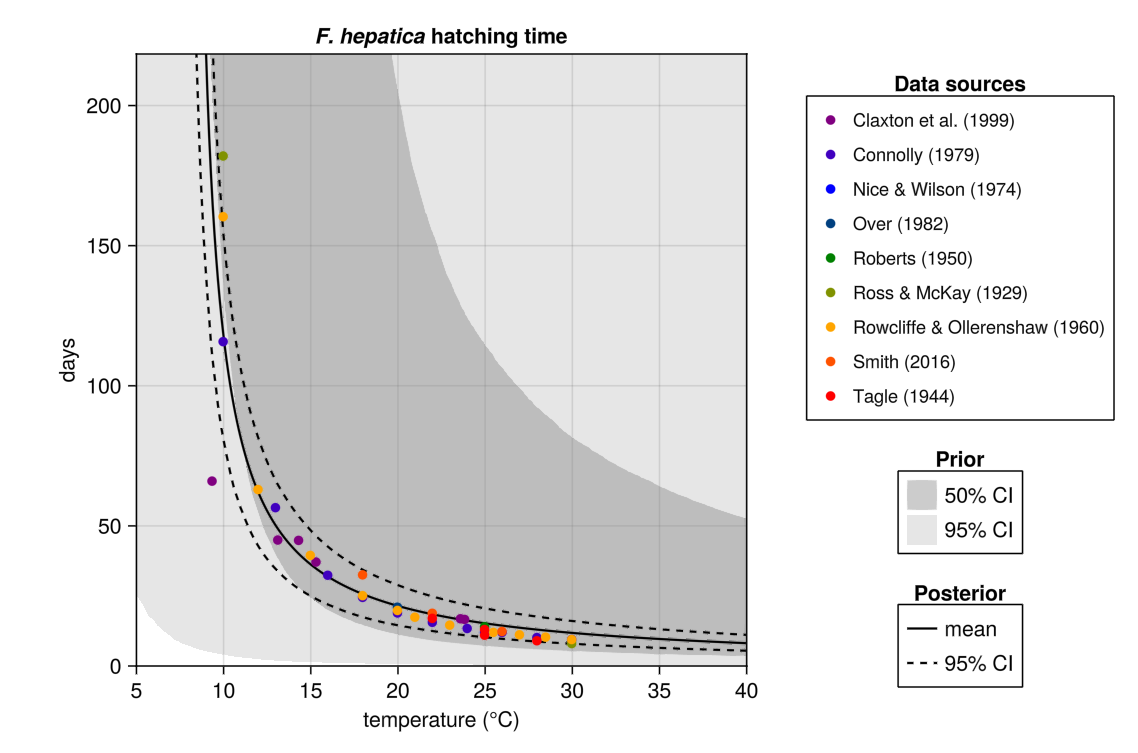

**Fig B. Data sources and prior and posterior distributions for *F. hepatica* hatching time**.

**Table B: Prior and posterior distributions for hatching time**

|  | | | |
| --- | --- | --- | --- |
| **Parameter** | ***F. hepatica*** | ***F. gigantica*** | **Prior** |
| days | 263.0 (175.4-360.4) | 173.9 (107.0-236.3) | LogNormal(μ=6.0, σ=2.0) |
| Tmin | 7.78 (7.20-8.27) | 13.9 (13.5-14.3) | Normal(μ=10.0, σ=5.0) |

#### 1.2 Prepatent period

Prepatent period is the time required from infection to the onset of shedding. As for the hatching time, we used a growing degree day model to fit data, such that:

$$PP\left( T \right)= \left\{ \begin{aligned} \frac{days}{T-Tmin}, &T>Tmin \\ 0, &T \leq Tmin \end{aligned} \right.$$

Where *PP* is the prepatent period, *days* is the number of growing degree days, and *Tmin* is the minimal temperature required for development. *Days* was again a random effect and *Tmin* assumed to be constant between studies.

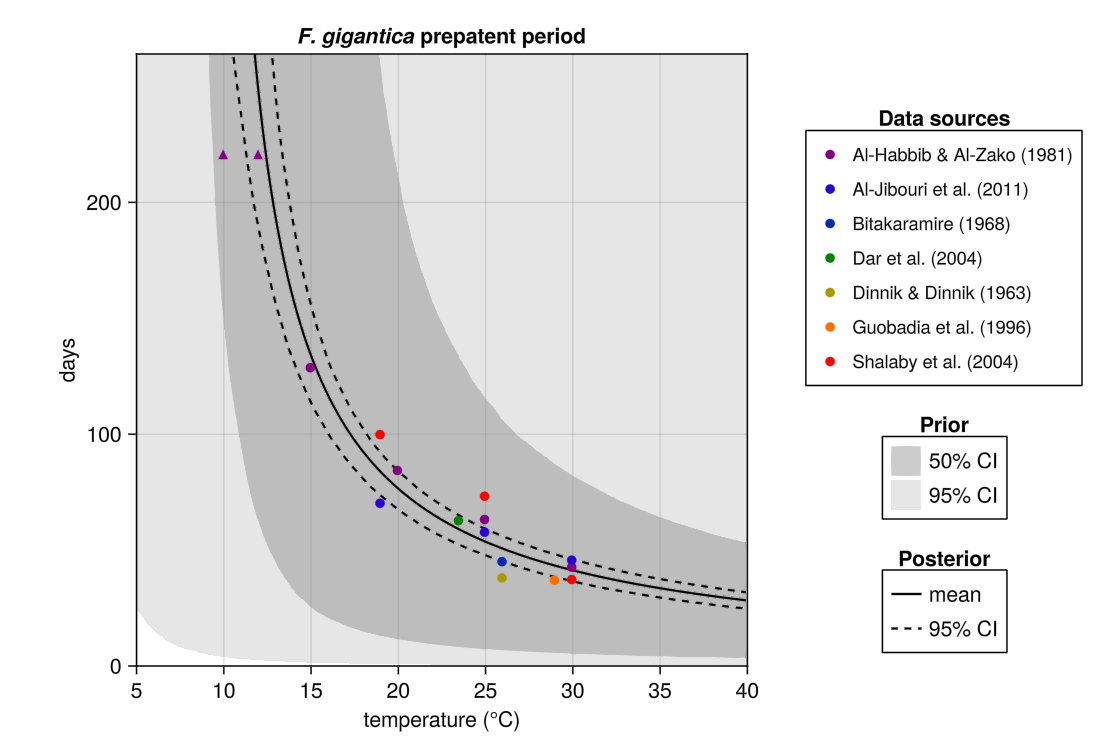

**Fig C. Data sources and prior and posterior distributions for *F. gigantica* prepatent period**.

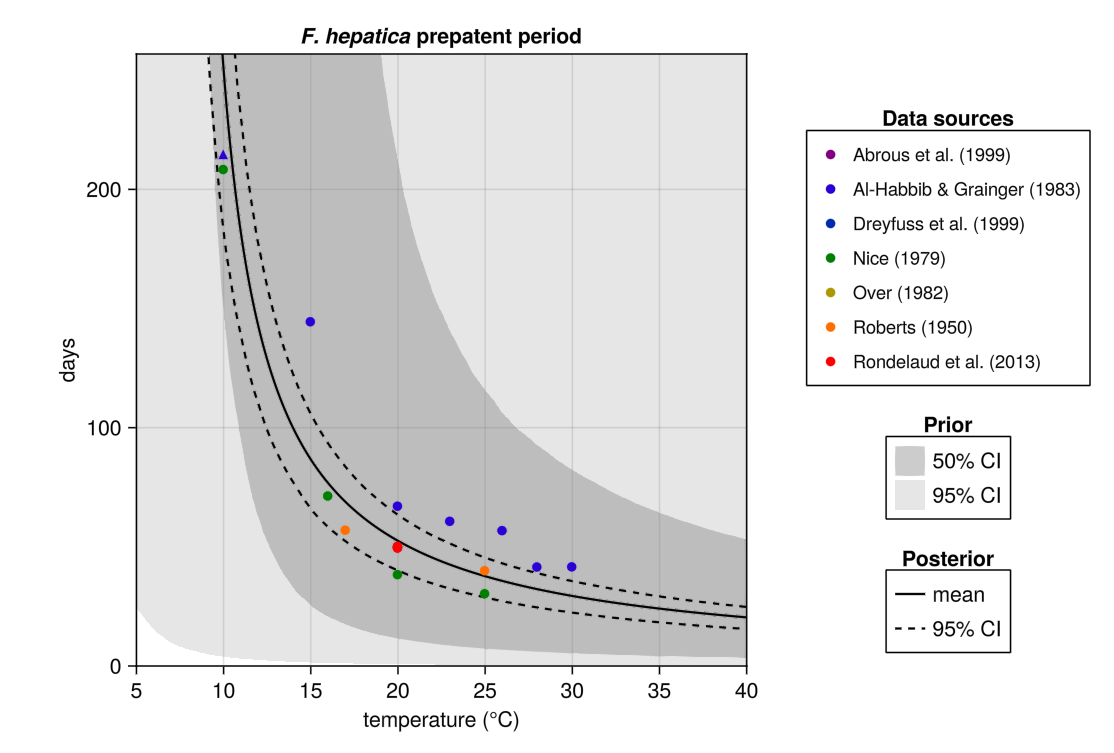

**Fig D. Data sources and prior and posterior distributions for *F. hepatica* prepatent period**.

**Table C: Prior and posterior distributions for prepatent period**

|  | | | |
| --- | --- | --- | --- |
| **Parameter** | ***F. hepatica*** | ***F. gigantica*** | **Prior** |
| days | 667.7 (508.9-823.3) | 898.9 (759.4-1037.8) | LogNormal(μ=6.0, σ=2.0) |
| Tmin | 7.33 (6.51-7.97) | 8.30 (7.05-9.78) | Normal(μ=10.0, σ=5.0) |

#### 1.3 Infection efficiency

Infection efficiency is the chance that a miracidium successfully infects a snail. We modelled this as a Gaussian function, such that:

$$IE\left( T \right)=rmax*e^{\frac{{-(T-Topt)}^{2}}{{2\sigma}^{2}}}$$

Where *IE* is the infection efficiency, *Topt* is the optimal temperature, *rmax* is the infection efficiency at *Topt* (bounded to between 0 and 1), and *σ* dictates the slope of the curve.

For *F. hepatia,* Christensen et al (1976) used radioactive metacercaria to estimate infection efficiency of *F. hepatica*, whereas other studies counted the number of infected snails. We therefore used separate priors for *rmax* for this study, that did not influence the final *rmax* estimate. We used a Binomial distribution for studies with counts for infected snails.

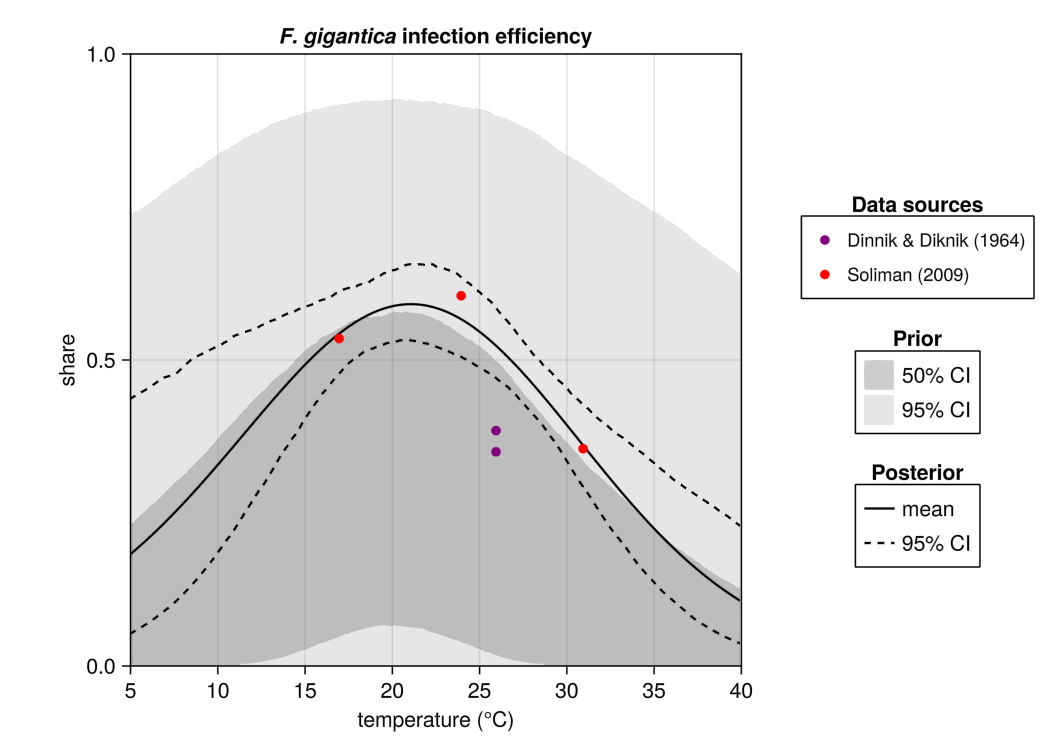

**Fig E. Data sources and prior and posterior distributions for *F. gigantica* infection efficiency**.

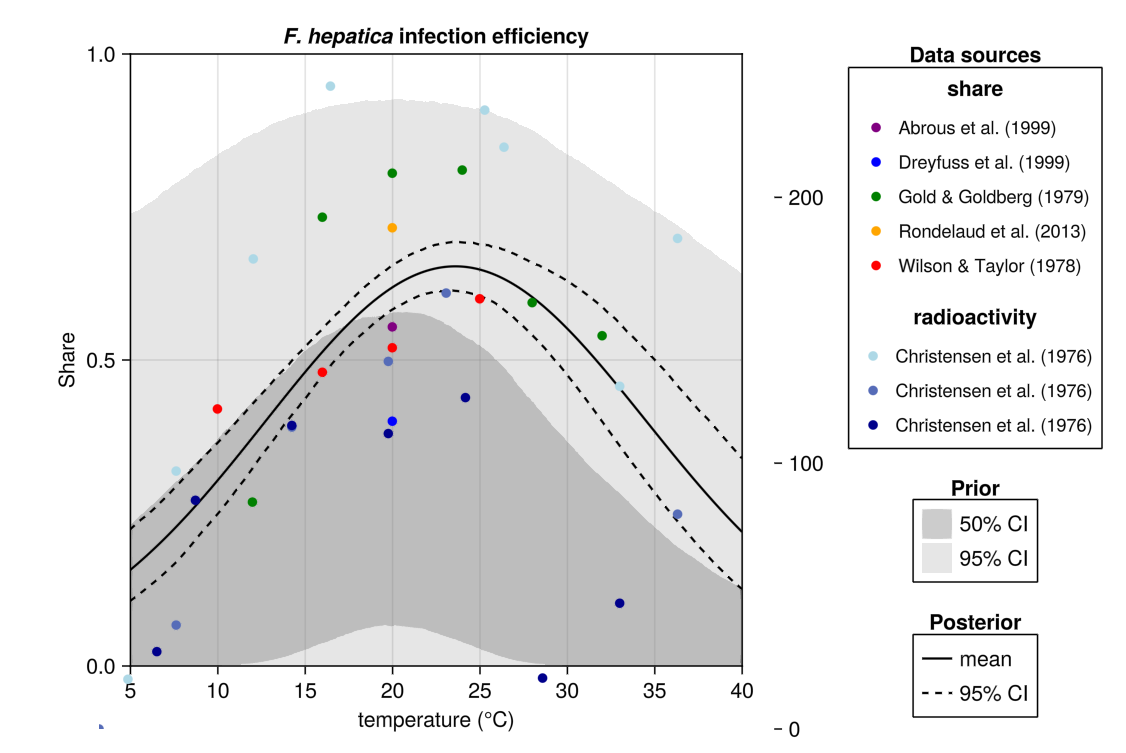

**Fig F. Data sources and prior and posterior distributions for *F. hepatica* infection efficiency.**

**Table D: Prior and posterior distributions for infection efficiency**

|  | | | |
| --- | --- | --- | --- |
| **Parameter** | ***F. hepatica*** | ***F. gigantica*** | **Prior** |
| rmax | 0.606 (0.424-0.753) | 0.513 (0.216-0.769) | Uniform(a=0.0, b=1.0) |
| Topt | 22.5 (21.5-23.5) | 21.2 (18.3-22.9) | Normal(μ=20.0, σ=5.0) |
| σ | 9.68 (8.62-11.01) | 9.82 (7.49-15.32) | Exponential(θ=10.0) |

#### 1.4 Cercarial release

For cercarial release, we also used a Gaussian curve, but log-transformed the observations before fitting, such that:

$$log(CR\left( T \right))=rmax*e^{\frac{{-(T-Topt)}^{2}}{{2\sigma}^{2}}}$$

Where *CR* is the cercarial release, *Topt* is the optimal temperature, *rmax* is the infection efficiency at *Topt* (bounded to between 0 and 1), and *σ* dictates the slope of the curve.

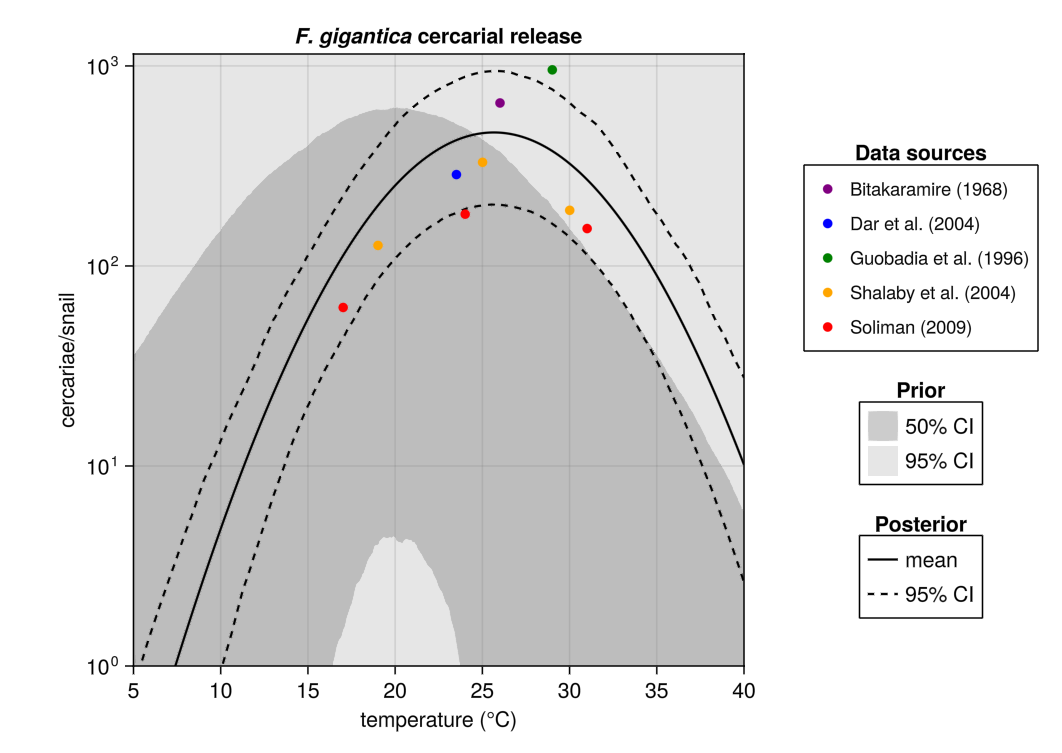

**Fig G. Data sources and prior and posterior distributions for *F. gigantica* cercarial release**.

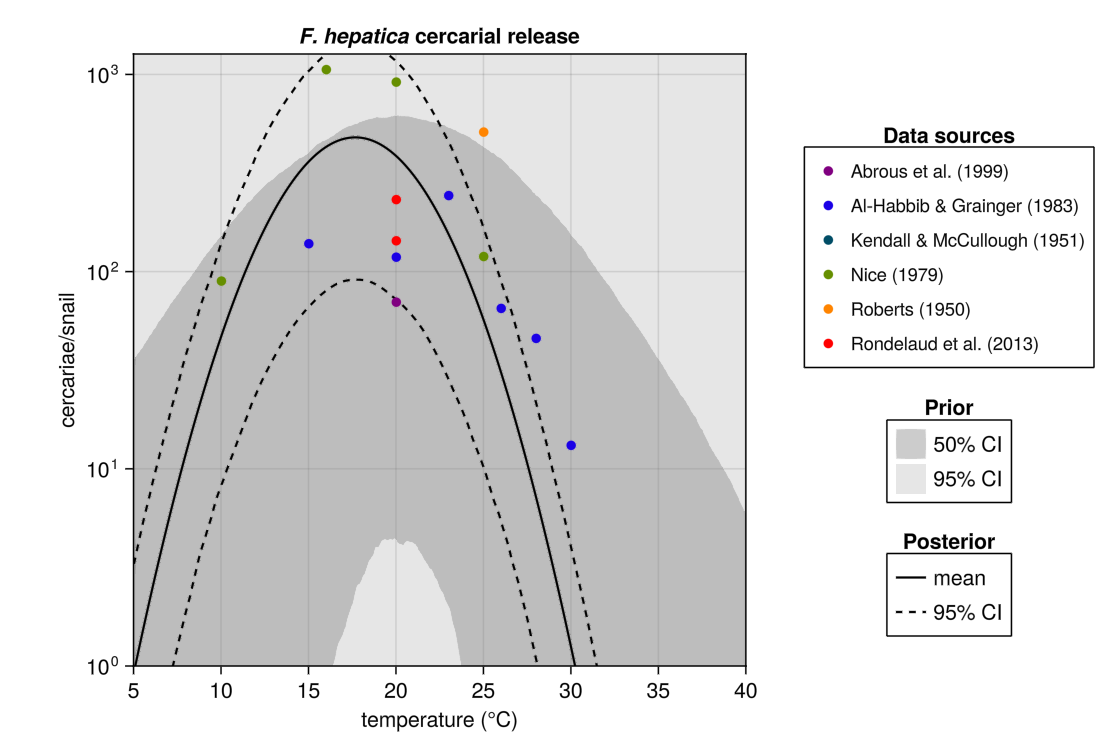

**Fig H. Data sources and prior and posterior distributions for *F. hepatica* cercarial release**.

**Table E: Prior and posterior distributions for cercarial release**

|  | | | |
| --- | --- | --- | --- |
| **Parameter** | ***F. hepatica*** | ***F. gigantica*** | **Prior** |
| rmax | 479.8 (91.1-1445.9) | 465.4 (202.7-943.1) | Normal(μ=6.0, σ=2.0) |
| Topt | 17.7 (17.4-17.9) | 25.6 (25.2-26.1) | Normal(μ=20.0, σ=5.0) |
| σ | 3.56 (3.25-3.86) | 5.14 (4.65-5.79) | Exponential(θ=1.0) |

#### 1.5 Hatching success

Hatching success is the number of eggs that successfully hatch. This was modelled as a function with three parts: it is zero beyond some critical minimum temperature, then follows a logistic function to decline towards zero.

$$HS\left( T \right)= \left\{ \begin{aligned} 0 , T<T_{min} \\ 1+\frac{r_{Tmax}-1}{T_{max}-T_{min}}*(T-T_{\min}), T_{min}\leq T\leq T_{max} \\ 2*r_{T_{max}}*\frac{e^{k*\left( T_{max}-T \right)}}{{1+ e}^{k*\left( T_{max}-T \right)}}, T >T_{max} \end{aligned} \right.$$

Here *HS* is the hatching success, *Tmin* is the temperature below which hatching success is 0, *Tmax* is the temperature above which hatching success declines rapidly, and *k* determines how quickly hatching success drops beyond *Tmax*.

All studies reported hatching success as the number of successfully hatched eggs, and we assumed a Binomial distribution.

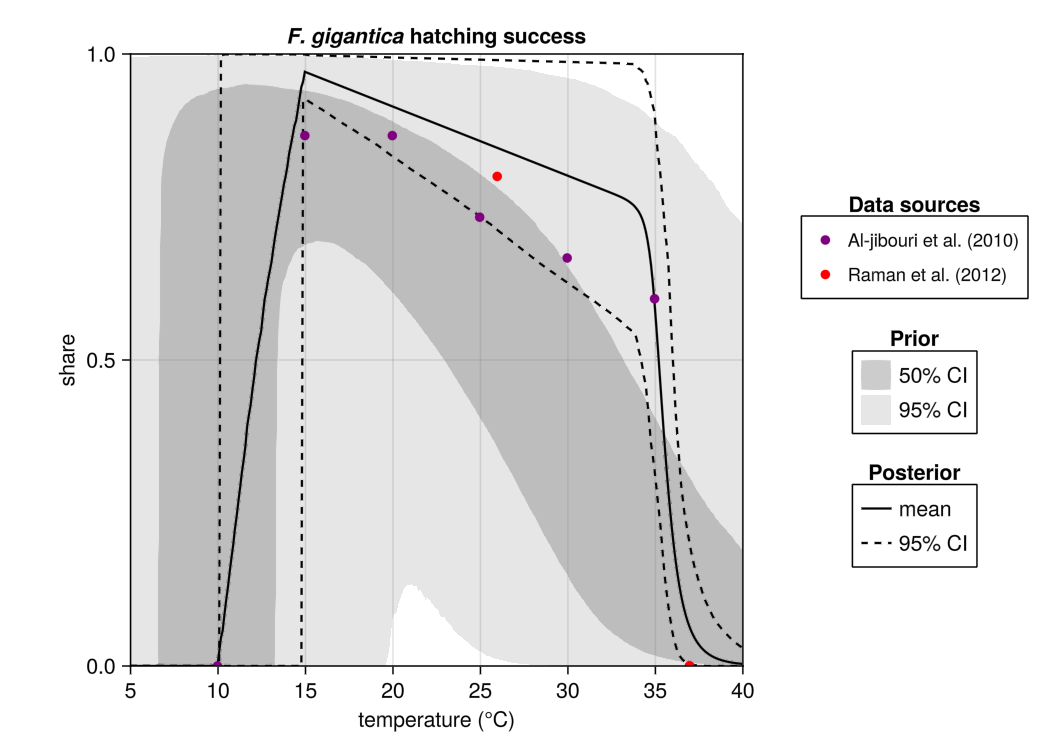

**Fig I. Data sources and prior and posterior distributions for *F. gigantica* hatching success**

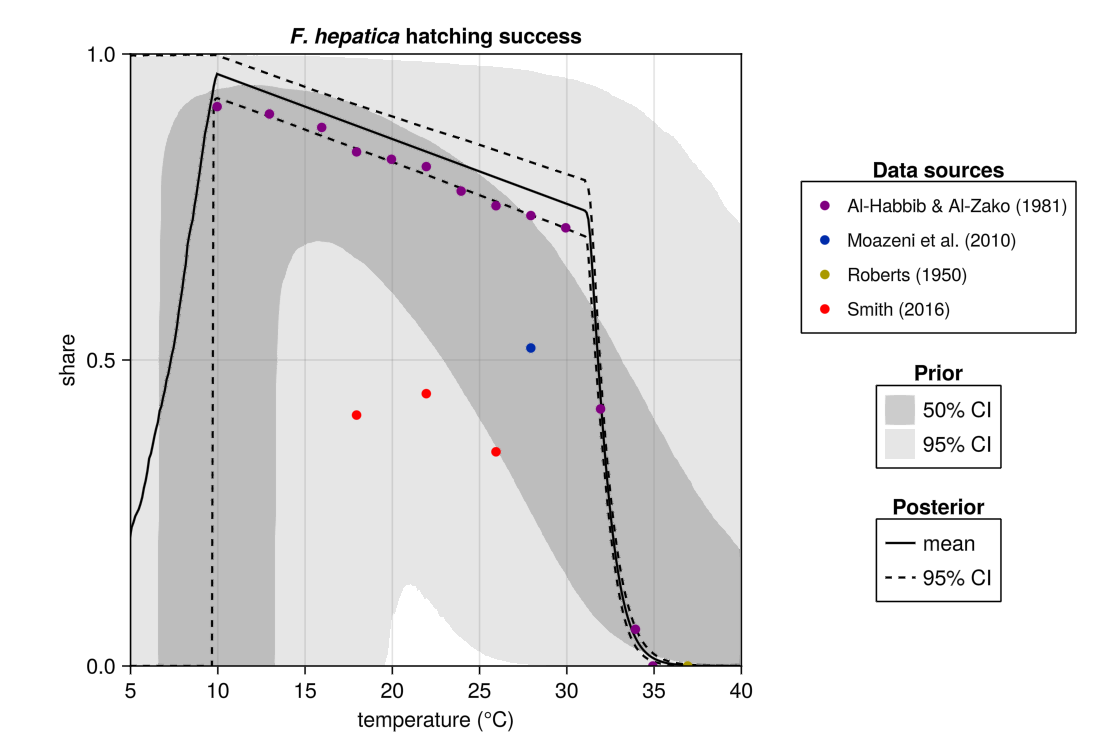

**Fig J. Data sources and prior and posterior distributions for *F. hepatica* hatching success**

**Table F. Prior and posterior distributions for hatching success**

|  | | | |
| --- | --- | --- | --- |
| **Parameter** | ***F. hepatica*** | ***F. gigantica*** | **Prior** |
| r_Tmax | 0.742 (0.700-0.793) | 0.747 (0.522-0.984) | Uniform(a=0.0, b=1.0) |
| k | 1.29 (1.11-1.47) | 1.67 (0.63-3.38) | Exponential(θ=0.5) |
| Tmin | 6.87 (2.75-9.79) | 12.3 (10.1-14.8) | Normal(μ=10.0, σ=5.0) |
| Tmax | 31.3 (31.1-31.5) | 34.7 (33.5-35.8) | Normal(μ=30.0, σ=5.0) |

### 2 Life cycle model sensitivity analysis

Beyond the life history traits described above, there are two life history traits in the life cycle model for which we did not fit traits but assumed constant. These are egg death rate and snail death rate. Fitting traits for these is difficult because of a paucity of data, and a very big spread in reported values between studies. In the field, mortality might be dominated by factors such as predation that are difficult to replicate in the lab. For our central estimate, we set both mortality rates to 0.01/day.

Fig K shows how the transmission strength index changes if these are assumed to be any combination of 0.002, 0.01, and 0.05. We find transmission strength is particularly sensitive to the snail death rate, with higher values for this parameter resulting in higher optimal temperatures.

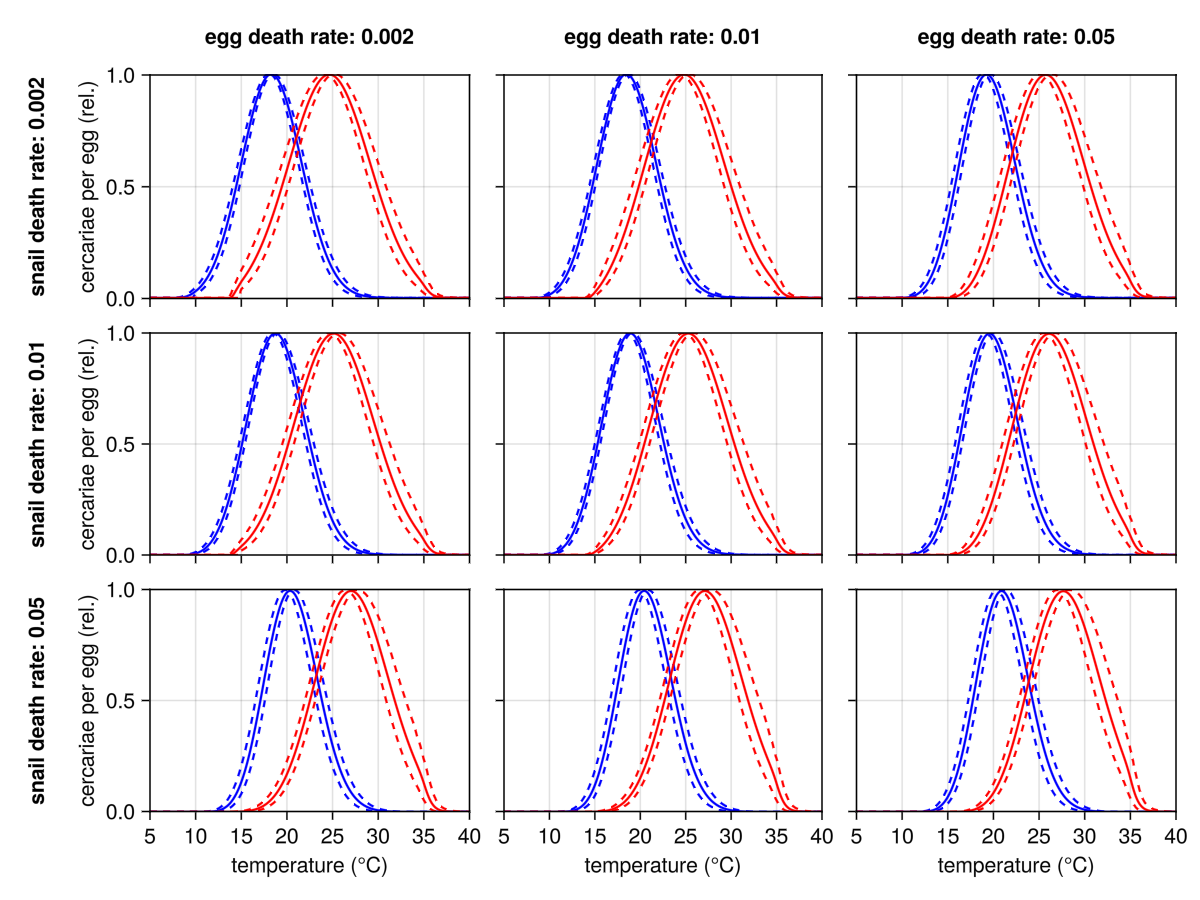

**Fig K**. **Sensitivity analysis for the effect of different snail and egg death rates on the transmission strength index.**

### 3 Species distribution modelling

#### 3.1 Literature search

PUBMED was searched for papers published between 2015 and 2025 using the following search query:

((gastropoda[mh] OR mollusca[mh] OR snails[mh]) OR

("Lymnaea natalensis" OR "L. natalensis" OR

"Lymnaea truncatula" OR "L. truncatula" OR

"Galba truncatula" OR "G. truncatula" OR

"Galba mweruensis" OR "G. mweruensis" OR

"Radix natalensis" OR "R. natalensis" OR

"Radix auricularia" OR "R. auricularia" OR

"Radix rubiginosa" OR "R. rubiginosa" OR

"Pseudosuccinea columella" OR "P. columella")

AND (Africa[MeSH Terms] OR Angola OR Benin OR Botswana OR Burkina OR Burundi OR Cameroon OR Central African Republic OR Chad OR Congo OR Cote d'Ivoire OR Djibouti OR Eswatini OR Eritrea OR Ethiopia OR Gabon OR Gambia OR Ghana OR Guinea OR Kenya OR Lesotho OR Liberia OR Libya OR Madagascar OR Malawi OR Mali OR Mauritania OR Mozambique OR Namibia OR Niger OR Nigeria OR Rwanda OR Senegal OR Sierra Leone OR Somalia OR South Africa OR Sudan OR Swaziland OR Tanzania OR Togo OR Tunisia OR Uganda OR Zambia OR Zaire OR Zimbabwe)

PUBMED articles that were then screened manually for relevance. Relevant articles included those that reported snail presence of at least one species of interest for this study, along with location information so the data could be georeferenced. Any articles written in other languages other than English, or that were not in SSA were excluded. Once a paper was determined relevant, data on snail species and location were manually extracted. A second random reviewer validated the extraction by sampling 25% of the relevant papers to re-do extraction to validate extraction was completed accurately. Authors were contacted for missing data or when disaggregated data were required for relevant papers.

#### 3.2 Occurrence records

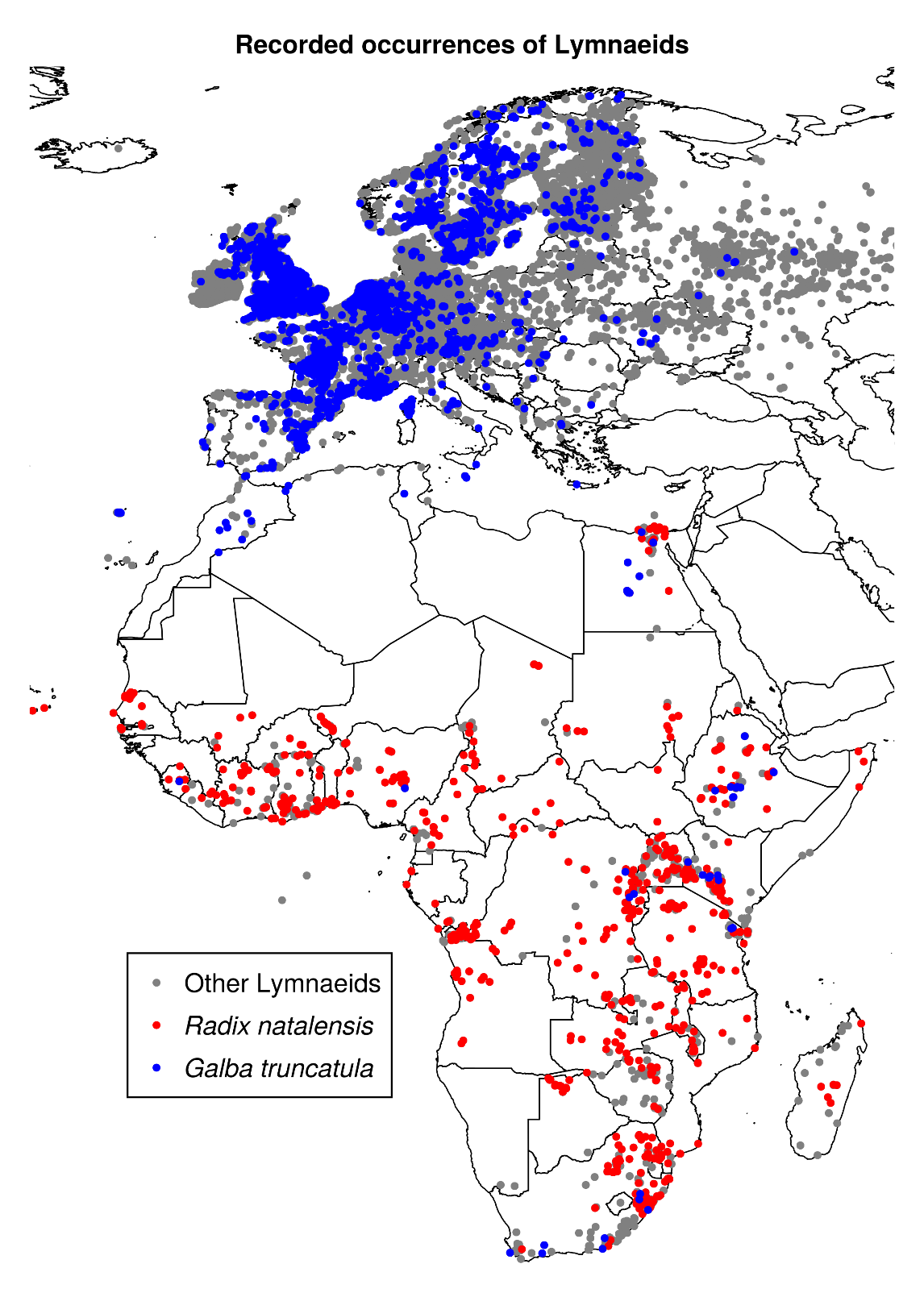

**Fig L. Occurrence records of *Radix natalensis*, *Galba truncatula*, and other lymnaeid snails.** Other lymnaeid occurrences were used as background points for species distribution modelling.

### 4 Projections

Fig M shows projected future *Fasciola* transmission risk under the SSP126 scenario. Figs N-O show projected transmission strength under the SSP126 and SSP370 scenarios. Figs P-Q show projected host suitability under the SSP126 and SSP370 scenarios. Finally, Figs R-S show projected transmission risk under the SSP370 scenario, for each hydrological and global circulation model used and for *F. hepatica* and *F. gigantica*, respectively.

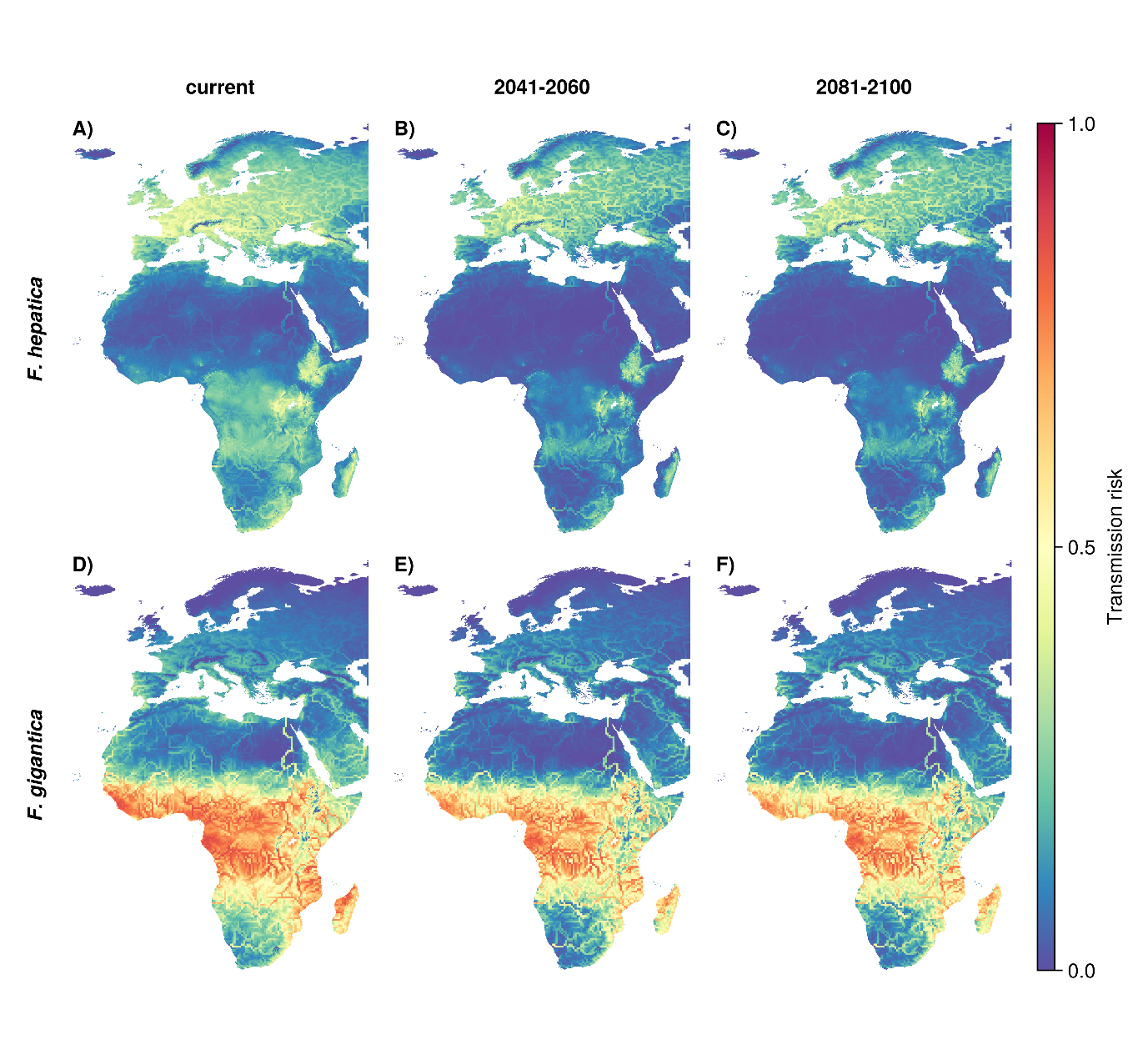

**Fig M. Projected fascioliasis transmission risk under the SSP126 scenario with low emissions.**

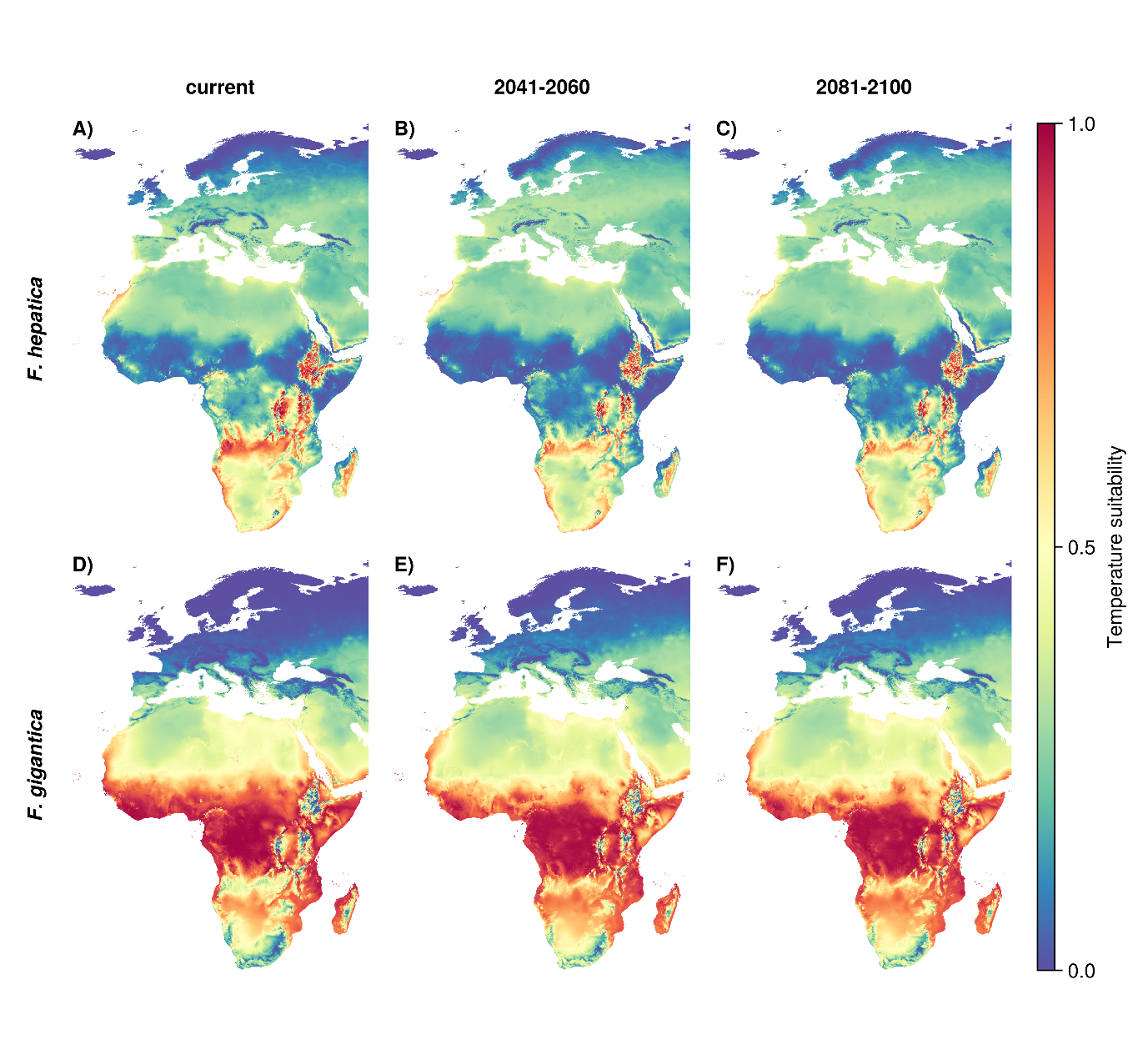

**Figure N. Projected temperature-dependent transmission potential under the SSP126 scenario with low emissions.**

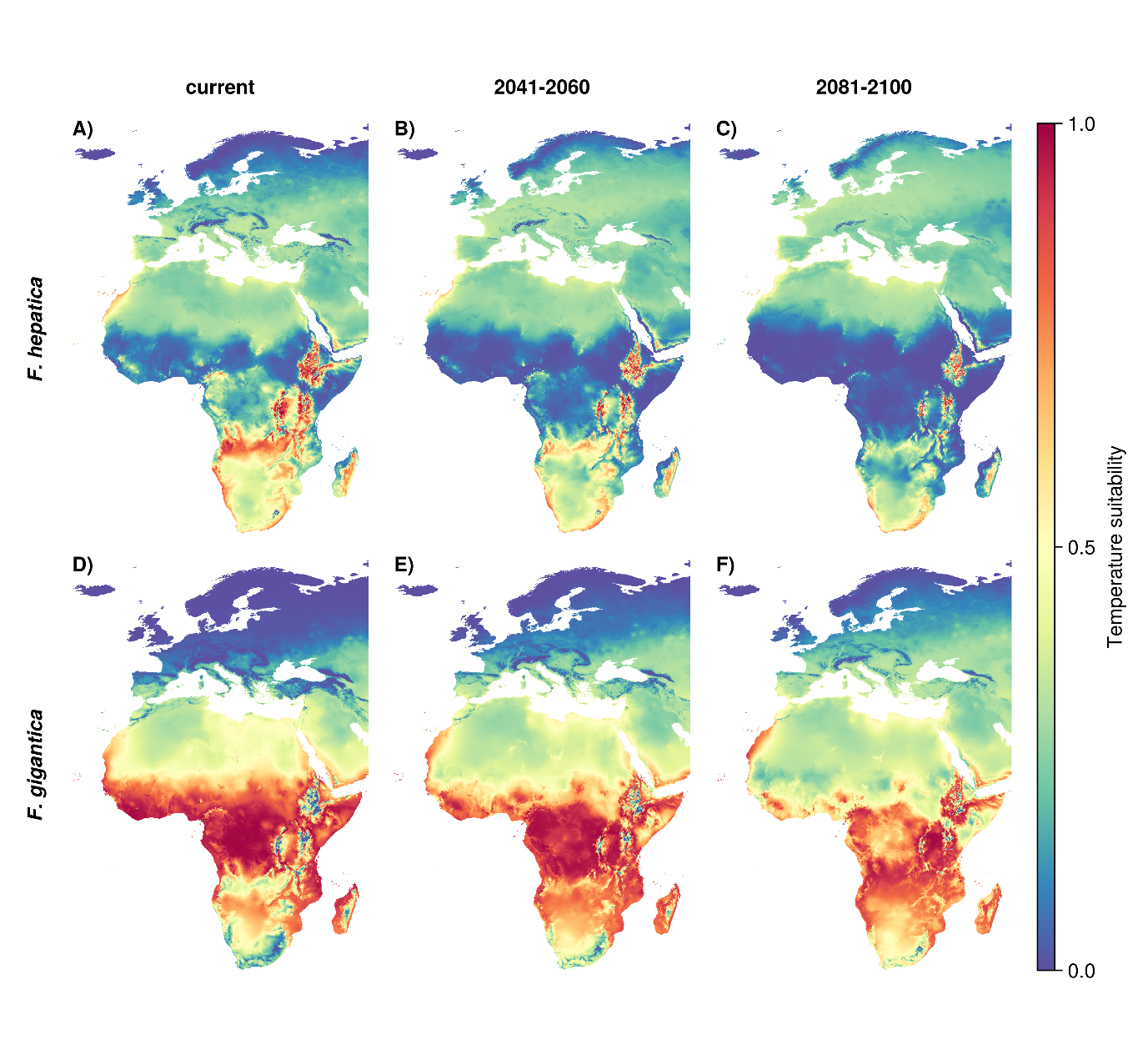

**Figure O. Projected temperature-dependent transmission potential under the SSP370 scenario with high emissions.**

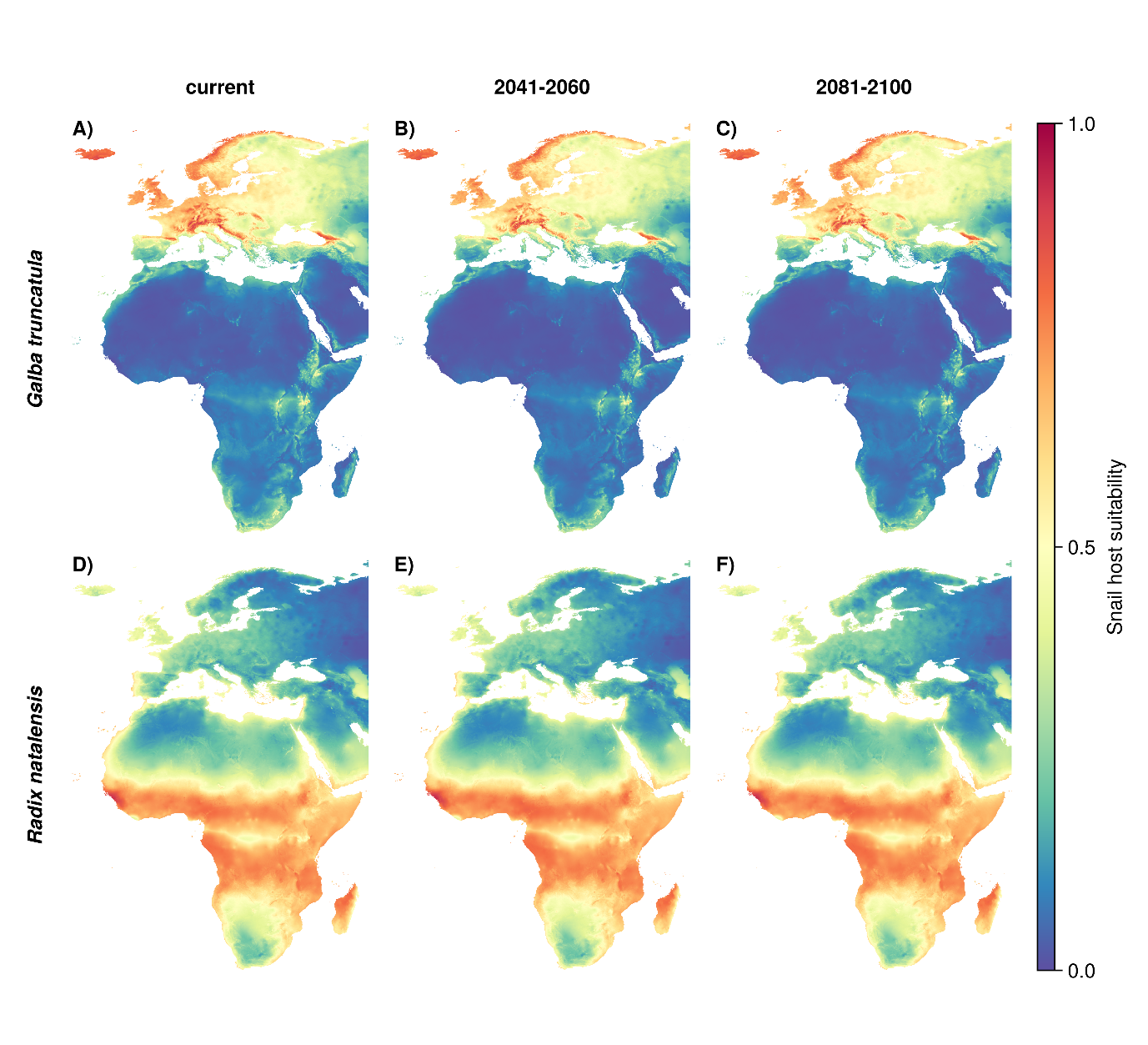

**Fig P. Projected snail host suitability under the SSP126 scenario with low emissions.**

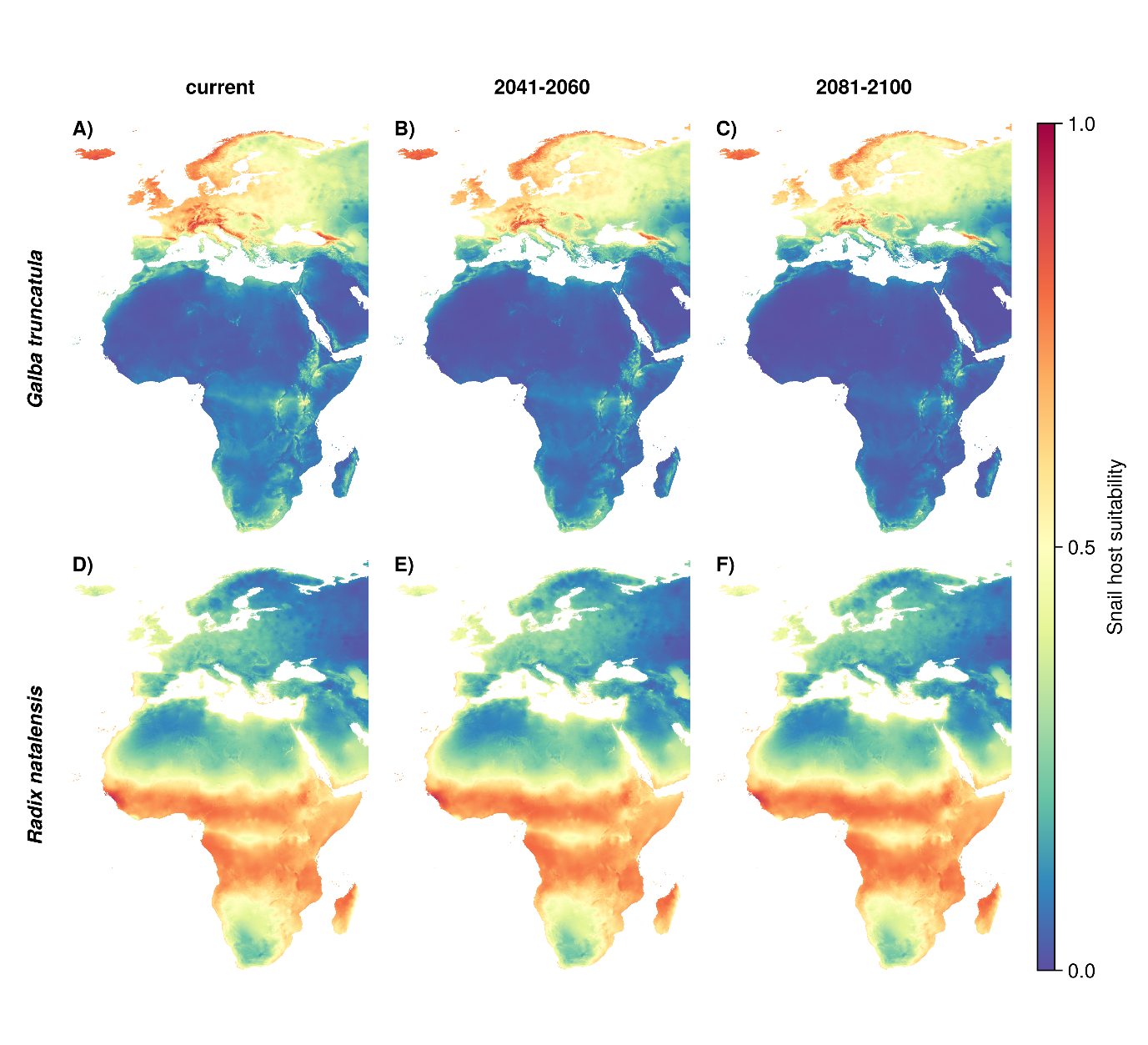

**Fig Q. Projected snail host suitability under the SSP370 scenario with high emissions.**

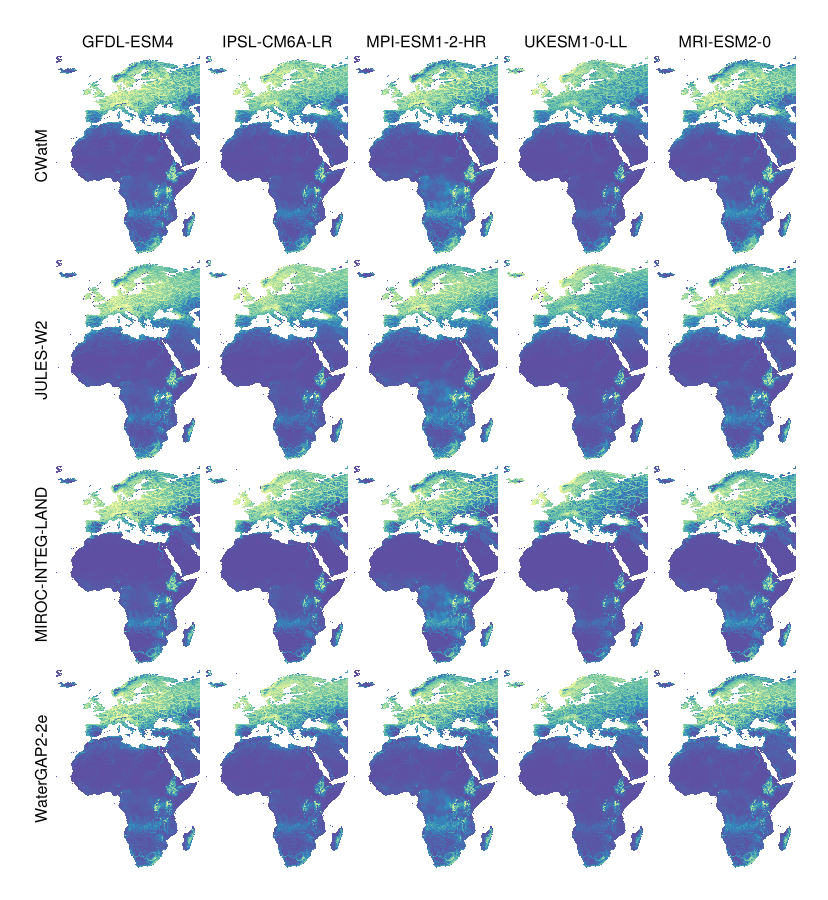

**Fig R. Projected transmission risk for *F. hepatica* under each combination of global hydrological model (rows) and global circulation model (columns).** UKESM1-0-LL is known to predict warming beyond the plausible range and was therefore excluded from the mean estimates presented in the paper.

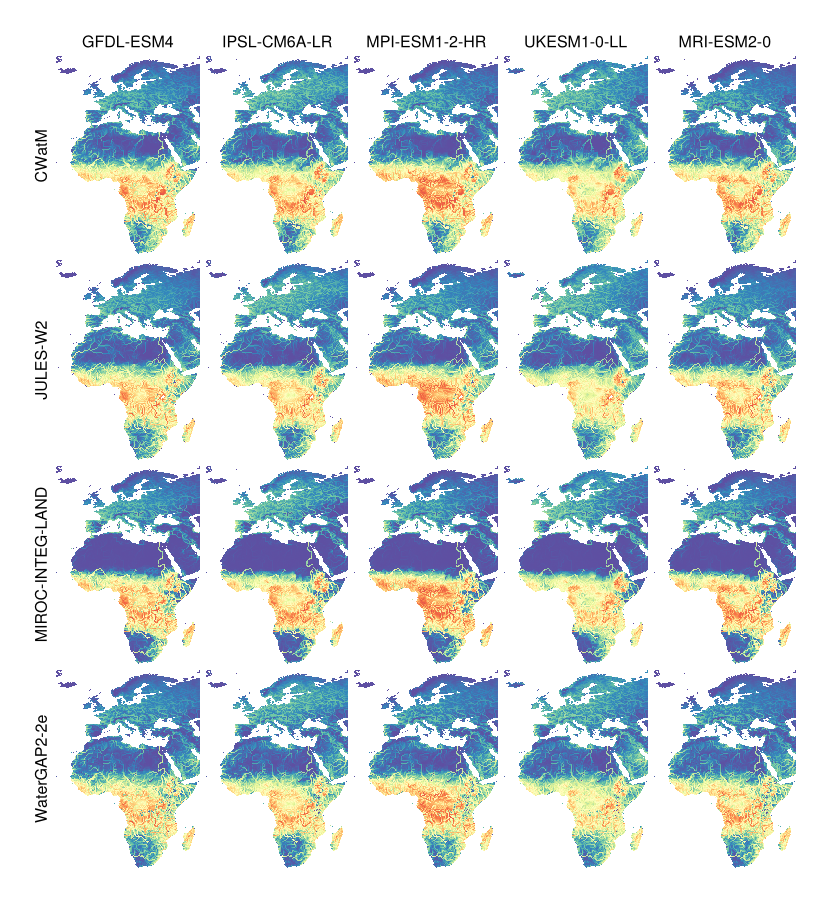

**Fig S. Projected transmission risk for *F. gigantica* under each combination of global hydrological model (rows) and global circulation model (columns).**

Nice, Nigel George. ‘Aspects of the Biology of Fasciola Hepatica and Its Intermediate Snail Host Lymnaea Truncatula’. PhD Thesis, University of York, 1979.

Over, Hans J. ‘Ecological Basis of Parasite Control: Trematodes with Special Reference to Fascioliasis’. *Veterinary Parasitology* 11, no. 1 (1 August 1982): 85–97. <https://doi.org/10.1016/0304-4017(82)90123-6>.

Phalee, Anawat, Chalobol Wongsawad, Amnat Rojanapaibul, and Jong-Yil Chai. ‘Experimental Life History and Biological Characteristics of Fasciola Gigantica (Digenea: Fasciolidae)’. *The Korean Journal of Parasitology* 53, no. 1 (February 2015): 59–64. <https://doi.org/10.3347/kjp.2015.53.1.59>.

Raman, M, R Selvarani N Jeyathilakan, C Soundararajan, Asha Alex, and G Ravikumar. ‘Maintenance of Radix Snails and Artificial Infection with Fasciola Gigantica Miracidium in Laboratory Condition’, 2012.

Rao, M. P. Chandra. ‘On the Comparative Susceptibility of Lymnaea Natalensis (Kraus) and L. Rufescens (Gray) to Infection with Fasciola Gigantica (West African Strain) and the Tissue Responses in the Snails’. *Journal of Helminthology* 40, no. 1–2 (June 1966): 131–40. <https://doi.org/10.1017/S0022149X00034143>.

Roberts, Enid W. ‘Studies on the Life-Cycle of Fasciola Hepatica (Linnaeus) and of Its Snail Host, Limnaea (Galba) Truncatula (Müller), in the Field and Under Controlled Conditions in the Laboratory’. *Annals of Tropical Medicine & Parasitology* 44, no. 2 (1 July 1950): 187–206. <https://doi.org/10.1080/00034983.1950.11685441>.

Rondelaud, Daniel, Amal Titi, Philippe Vignoles, Abdeslam Mekroud, and Gilles Dreyfuss. ‘Consequence of Temperature Changes on Cercarial Shedding from Galba Truncatula Infected with Fasciola Hepatica or Paramphistomum Daubneyi’. *Parasite* 20 (2013): 10. <https://doi.org/10.1051/parasite/2013009>.

Ross, Sir Ian Clunies, and AC McKay. ‘The Bionomics of Fasciola Hepatica in New South Wales and of the Intermediate Host Limnea Brazieri (Smith)’. *Bulletin (Council for Scientific and Industrial Research (Australia))* 43 (1929).

Rowcliffe, S. A., and C. B. Ollerenshaw. ‘Observations on the Bionomics of the Egg of Fasciola Hepatica’. *Annals of Tropical Medicine & Parasitology* 54, no. 2 (June 1960): 172–81. <https://doi.org/10.1080/00034983.1960.11685973>.

Shalaby, Ismail, Mohamed Hassan, Maha Soliman, and N. Shrief. ‘Factors Affecting Dynamics of Metacercarial Productivity of Fasciola Gigantica from Its Snail Host’. *Pakis. J. Biol. Sci.* 7 (1 March 2004). <https://doi.org/10.3923/pjbs.2004.393.398>.

Sharma, R. L., The late D. N. Dhar, and O. K. Raina. ‘Studies on the Prevalence and Laboratory Transmission of Fascioliasis in Animals in the Kashmir Valley’. *British Veterinary Journal* 145, no. 1 (1 January 1989): 57–61. <https://doi.org/10.1016/0007-1935(89)90010-9>.

Smith, Daniel Barnaby. ‘Predicting Temporal Changes in Fasciola Hepatica Abundance from Climatic Variables’, 2016.

Soliman, Maha F. M. ‘*Fasciola Gigantica*: Cercarial Shedding Pattern from *Lymnaea Natalensis* after Long-Term Exposure to Cadmium at Different Temperatures’. *Experimental Parasitology* 121, no. 4 (1 April 2009): 307–11. <https://doi.org/10.1016/j.exppara.2008.12.001>.

Tagle, I. ‘Observaciones Sobre La Evolución de La Fasciola Hepatica Linneo 1758. Comprobación Del Huésped Intermediario En Chile’ 46, no. 47 (1944): 232.

Wilson, R. A., and Susan L. Taylor. ‘The Effect of Variations in Host and Parasite Density on the Level of Parasitization of Lymnaea Truncatula by Fasciola Hepatica’. *Parasitology* 76, no. 1 (February 1978): 91–98. <https://doi.org/10.1017/S0031182000047429>.

1. Department of Veterinary and Animal Sciences, University of Copenhagen, Copenhagen, Denmark
2. Center for Macroecology, Evolution, and Climate, Globe Institute, University of Copenhagen, Copenhagen, Denmark
3. School of Agriculture and Science, College of Agriculture, Engineering and Science, University of KwaZulu-Natal, Durban, South-Africa
4. One Health Center for Zoonoses and Tropical Veterinary Medicine, Ross University School of Veterinary Medicine, Basseterre, Saint Kitts and Nevis
5. School of Medicine, College of Health Sciences, University of KwaZulu-Natal, Durban, South Africa
6. Department of Physics, Geography and Environmental Sciences, School of Natural Sciences, Great Zimbabwe University, Masvingo, Zimbabwe
7. Geosciences Department, School Geosciences, Disaster and Sustainable Development, Faculty of Science, Bindura University of Science and Education, Bindura, Zimbabwe
8. National Institute for Medical Research (NIMR), Mwanza Centre, Mwanza, Tanzania
9. Swiss Tropical and Public Health Institute, Allschwil, Switzerland
